# A mechanism for the extension and unfolding of parallel telomeric G-quadruplexes by human telomerase at single-molecule resolution

**DOI:** 10.1101/2020.02.26.965269

**Authors:** Bishnu P. Paudel, Aaron Lavel Moye, Hala Abou Assi, Roberto El-Khoury, Scott B. Cohen, Monica L. Birrento, Siritron Samosorn, Kamthorn Intharapichai, Christopher G. Tomlinson, Marie-Paule Teulade-Fichou, Carlos González, Jennifer L. Beck, Masad J. Damha, Antoine M. van Oijen, Tracy M. Bryan

## Abstract

Telomeric G-quadruplexes (G4) were long believed to form a protective structure at telomeres, preventing their extension by the ribonucleoprotein telomerase. Contrary to this belief, we have previously demonstrated that parallel-stranded conformations of telomeric G4 can be extended by human and ciliate telomerase. However, a mechanistic understanding of the interaction of telomerase with structured DNA remained elusive. Here, we use single-molecule fluorescence resonance energy transfer (smFRET) microscopy and bulk-phase enzymology to propose a mechanism for the resolution and extension of parallel G4 by telomerase. Binding is initiated by the RNA template of telomerase interacting with the G-quadruplex; nucleotide addition then proceeds to the end of the RNA template. It is only through the large conformational change of translocation following synthesis that the G-quadruplex structure is completely unfolded to a linear product. Surprisingly, parallel G4 stabilization with either small molecule ligands or by chemical modification does not always inhibit G4 unfolding and extension by telomerase. These data reveal that telomerase is a parallel G-quadruplex resolvase.

---

Human chromosomes contain many guanine (G)-rich elements capable of forming four-stranded G-quadruplex (G4) structures (1–4). A planar G-quartet is formed when four Gs form hydrogen bonds in a cyclical manner via their Hoogsteen faces; such G-quartets stack on top of each other and form G-quadruplexes (5, 6). G-rich sequences are primarily located in promoter regions, intron and exon boundaries, origins of replication and telomeres (1–4). Vertebrate telomeres consist of many tandem repeats of the sequence TTAGGG, comprising a double-stranded region and a 3’ G-rich overhang (7). Fluorescence microscopy studies have shown the existence of G4 structures at the telomeric regions of fixed human cells using G4-specific antibodies or stabilizing ligands (8, 9). *In vitro*, telomeric DNA can adopt many different conformations of G4, including inter- or intramolecular forms, arranged in parallel or anti-parallel orientation depending upon the directionality of the DNA backbone (10). Conditions that may mimic those in the nucleus, including high DNA concentration and water depletion, favour parallel conformations of human telomeric G4 (11–13). However, the *in vivo* conformation(s) and biological significance of telomeric G4 in human cells remain elusive and undetermined.

Telomere shortening occurs with every cell division in normal human somatic cells, since conventional DNA polymerases cannot completely synthesize the telomeric end (14–16). Telomerase, a telomere-specific ribonucleoprotein enzyme complex, extends telomeric DNA in cancer cells, stem cells and cells of the germline, using a unique mechanism of processive rounds of reverse transcription (17–22). Human telomerase minimally consists of the highly conserved telomerase reverse transcriptase protein (hTERT) and an RNA component containing an 11 nucleotide (nt) template sequence (hTR) (23). Telomerase extends its telomeric substrate by first binding to telomeric DNA, followed by hybridization of the RNA template with the DNA. Second, telomerase extends the substrate DNA by using its RNA as a template for nucleotide addition to the DNA 3’ end. Once the template 5’ boundary has been reached, telomerase translocates downstream, resulting in re-alignment of the RNA template with the new 3’ end of the product DNA (19, 22).

One function of telomeric G4 may be to act as a ‘cap’, protecting the telomere from DNA degradation (24). It has been widely believed that G4 formation within the 3’ telomeric overhang blocks telomere extension by telomerase, since early *in vitro* studies had shown that G4 structures inhibit telomerase extension of a telomeric DNA substrate (25, 26). In addition, stabilization of G4 with small-molecule ligands has been shown to more effectively inhibit telomerase activity, suggesting that chemical stabilization of G4 structures may be a viable anti-cancer therapeutic strategy (27–29). However, the above-mentioned studies did not distinguish between G4 conformations, and used oligonucleotides that likely folded into anti-parallel or ‘hybrid’ G4 forms as a telomerase substrate. A variety of helicases, such as RECQ5, WRN, BLM, FANC-J and RHAU, recognize and resolve G4 in a conformation-specific manner (30–33). Similarly, we have demonstrated that telomerase can bind and extend specific conformations of telomeric G4 - those that are parallel-stranded and intermolecular - and that this property of telomerase is well conserved from ciliates to human (34–36). This suggests that if telomeric G-quadruplexes were to adopt a parallel conformation *in vivo*, they may indeed be extended by telomerase, contrary to what had been previously hypothesized.

Telomerase extension reactions in the presence of individual nucleotides demonstrated that the 3’ end of a G4 substrate aligns correctly with the RNA template of telomerase, suggesting that at least partial resolution of G4 structure is required for telomerase extension (34). However, the evidence for partial disruption of the G4 was indirect, and a mechanistic understanding of telomerase resolution of G4 DNA, in the absence of a known helicase function of hTERT, remained elusive. Furthermore, since all previously-tested substrates were both parallel-stranded and intermolecular, it was unknown which of these properties form the basis of telomerase recognition. To directly study the mechanistic details of parallel G4 unfolding and extension by human telomerase, here we combined ensemble telomerase enzymatic assays and single-molecule FRET (smFRET) measurements *in vitro*, using both intramolecular and intermolecular parallel G4 as FRET sensors. We provide direct evidence that wild-type human telomerase can unfold both intra- and intermolecular parallel G4 completely in the presence of dNTPs, using a 3-step mechanism. First, telomerase binds the G4, partially changing its conformation in a process that involves binding of the hTR template sequence. Second, telomerase adds individual nucleotides to the 3’ end of the partially unfolded parallel G4. Lastly, the translocation of telomerase results in complete disruption of the G4 structure. Unexpectedly, stabilization of parallel intermolecular G4 using different G4 ligands did not inhibit telomerase unfolding and extension of this substrate. Overall, we provide a mechanistic explanation of conformation-specific telomerase extension of telomeric G4 at single-molecule resolution and demonstrate that small molecule-mediated inhibition of telomerase extension of telomeric G4 is topology-dependent.

## Results

### Human telomerase binds, unfolds and extends intramolecular parallel G-quadruplexes

Parallel, intermolecular G4 are substrates for telomerase, whereas intramolecular antiparallel or hybrid conformations are not (25, 26, 34, 35, 37). Determining whether it is the parallel or intermolecular nature of G4 structures that allows their recognition by telomerase has been difficult, since a 4-repeat human telomeric oligonucleotide does not readily fold into stable parallel intramolecular G4 at the concentrations used in *in vitro* assays, and instead exists as a mixture of topologies under most conditions (38–41). For this reason, we made use of the modified nucleotide 2’-fluoro-arabinoguanosine (2’F-araG), which induces parallel propeller-type G4 conformations (42). We have previously demonstrated that substitution of six guanosines in the telomeric sequence AGGG(TTAGGG)_3_ with 2’F-araG leads to a 15°C increase in *T*_m_ of the resulting intramolecular G4, and a shift from the usual antiparallel or hybrid topology of this sequence (11, 43) to a parallel conformation (Figure 1a) (44). Here, we demonstrate that G4 formed from the unmodified sequence (22G0) in KCl is a poor telomerase substrate, leading to the previously-observed stuttering pattern (26) in a direct telomerase extension assay involving incorporation of radiolabeled α^32^P-dGTP (Figure 1b). Substitution with 2’F-araG (22G3; see Supplementary Table 1 for sequence) restored the expected 6-nt repeat pattern of telomerase extension (Figure 1b), despite the increase in thermal stability of this G4. Thus, telomerase is able to extend parallel G4 structures, whether they are inter-or intramolecular.

**Figure 1:**
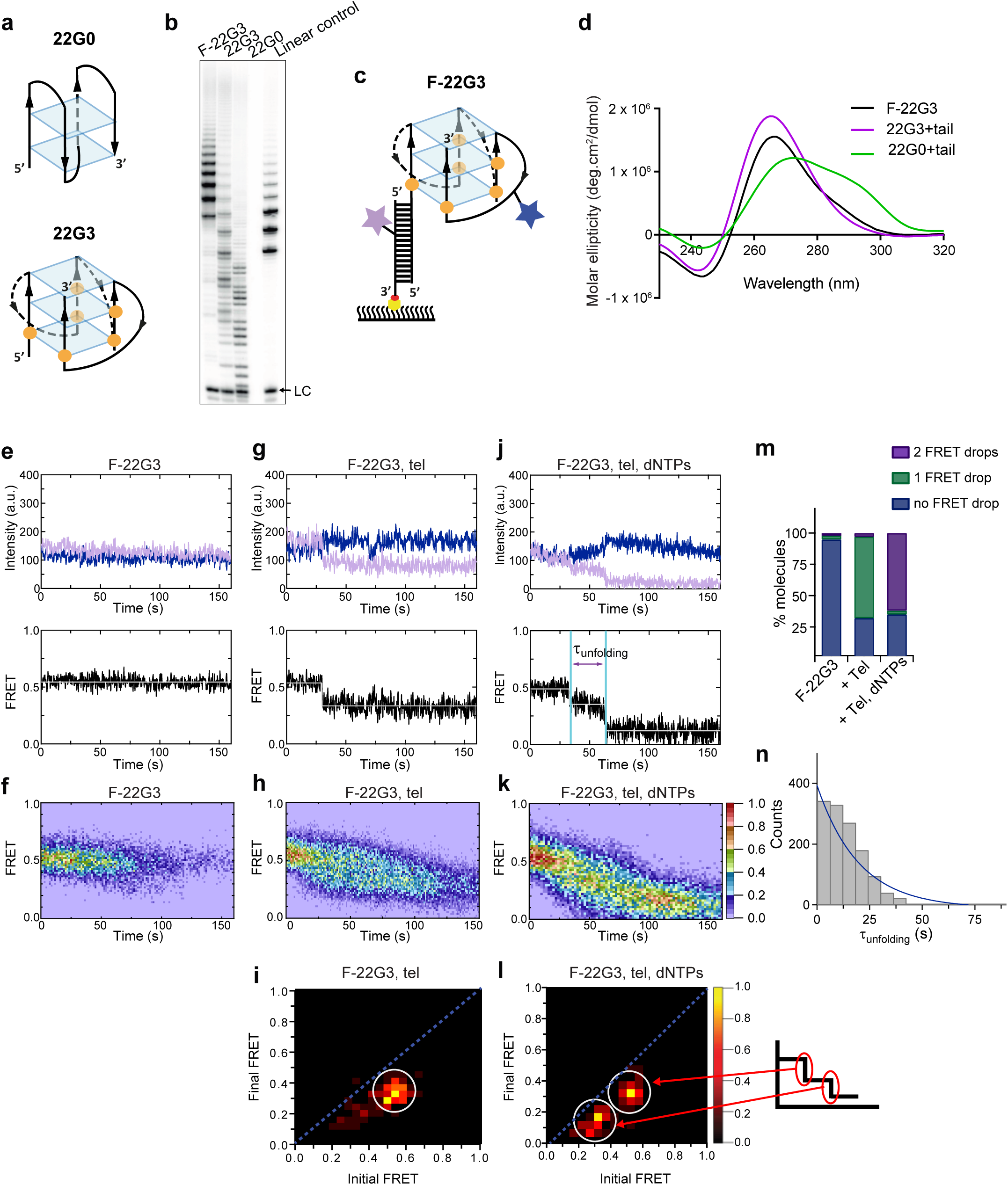
Telomerase unfolds and extends a parallel G-quadruplex. (**a**) Schematic of the likely topology of (**top**) an antiparallel G-quadruplex formed from unmodified telomeric 22-mer 22G0 and (**bottom**) a parallel unimolecular G-quadruplex formed from a telomeric 22-mer with 2’F-araG in the indicated positions (orange circles). (**b**) Telomerase extension assays in the presence of radiolabeled dGTP (α^32^P-dGTP). The extension products of 2 µM each of fluorescently-labeled 22G3 (F-22G3), unlabeled 22G3, unmodified 22G0 and linear control oligonucleotide Bio-L-18GGG (see Supplementary Table 1) were analyzed using denaturing polyacrylamide gel electrophoresis. LC: 5’-^32^P-labeled synthetic 12 nt DNA used as a recovery/loading control. (**c**) The F-22G3 FRET construct, showing the positions of the FRET dye pair and biotin for immobilization onto the functionalized coverslip. Blue star: AlexaFluor 555^TM^ (donor dye); purple star: AlexaFluor 647^TM^ (acceptor dye); orange circles: 2’F-araG; red circle: biotin; yellow circle: Neutravidin. (**d**) CD spectra of G4 (20 µM) formed from 2’F-araG-modified 5’-extended 22G3 labeled with a FRET dye (22G3+tail), and the same G4 after hybridization to a second oligonucleotide bearing a second FRET dye (F-22G3). A peak at 265 nm is characteristic of parallel G4; the slight shoulder at ∼290 nm in the F-22G3 spectrum is attributable to duplex DNA since it is absent from the spectrum of 22G3+tail. 22G0+tail is a control oligonucleotide of equal length and base composition to 22G3+tail, but with no 2’F-ara G substitutions or conjugated dye. (**e**),(**g**),(**j**) Representative single-molecule acceptor (purple) and donor (blue) intensities of F-22G3 molecules over time (top panels), and the FRET traces (bottom panels) representing the ratio of acceptor intensity to the sum of acceptor and donor intensities, of either F-22G3 alone (**e**), in the presence of telomerase (**g**), or in the presence of telomerase and dNTPs (**j**). (**f**),(**h**),(**k**) Heat maps of the distribution of FRET intensities over 0 - 150 s of F-22G3, either alone (n = 99) (**f**), in the presence of telomerase (n = 125) (**h**), or in the presence of telomerase and dNTPs (n = 90) (**k**). All plots include molecules collected in 4 – 6 independent experiments. For color key, see panel (**k**). (**i**),(**l**) TDPs showing the change in FRET value of molecules from experiments in (**h**) and (**k**) (n = 90 and 75, respectively). Schematic in (**l**) shows assignment of each TDP peak to one of the two steps of FRET reduction observed in single molecule traces. (**m**) Plot of the percentage of molecules showing no change in FRET, a single FRET drop, or a two-step FRET drop, in the experiments shown in (**e-l**). (**n**) The unfolding rate of F-22G3, calculated by fitting the dwell time distributions of the intermediate FRET state (see panel **j**) to a single exponential equation (n = 76 molecules).

To determine the mechanism of extension of intramolecular G4, we designed a version of 22G3 with a FRET donor dye (AlexaFluor 555™) on one of the propeller loops, and an extended 5’ tail; circular dichroism (CD) spectroscopy demonstrated that the G4 formed from this sequence retained a predominantly parallel topology (Supplementary Table 1 and Figure 1c,d; oligonucleotide 22G3+tail). The extended 5’ tail enabled hybridization with a second DNA oligonucleotide containing a FRET acceptor dye (AlexaFluor 647™) and a biotin for surface immobilization in smFRET studies; we refer to the assembled FRET-modified G4 as F-22G3 (Figure 1c). F-22G3 also retained a parallel G4 topology, with a slight shoulder at ∼290 nm attributable to the duplex portion of the molecule (Figure 1d). Assembled F-22G3 was efficiently extended by telomerase, and was possibly an even better substrate than 22G3, with the expected 6-nt repeat pattern (Figure 1b).

We predicted that this structure would yield a high FRET ratio as the donor and acceptor fluorophores would be in close proximity (Figure 1c), but when unfolded the strands would move apart and a low FRET ratio would be expected. We carried out smFRET experiments using the modified F-22G3 construct by tethering it on pegylated coverslips via a biotin-streptavidin-biotin linkage. Surface-immobilized G4s were excited using a 532 nm laser and signals emitted from both the donor and acceptor fluorophores were collected and their intensities measured over time (Figure 1e, top panel); both dyes provided a constant fluorescence signal over several minutes. The apparent FRET values between these two fluorophores were calculated by dividing the acceptor intensity by the sum of the donor and acceptor intensities (Figure 1e, bottom panel). Individual F-22G3 molecules all displayed a constant FRET ratio over time, indicating that they did not undergo any detectable conformational changes in the experimental time window. Using ∼100 such smFRET trajectories pooled from multiple independent experiments (sample sizes for each condition indicated in figure panels), a FRET heat map was constructed, showing the distribution of average FRET values (Figure 1f). The heat map showed that the mean FRET value (0.53 ± 0.05) remained unchanged over time; an alternative histogram representation of the same data grouped into 15 s bins confirms this conclusion (Supplementary Figure 1a). These data demonstrate that F-22G3 is stable over time.

Next, we tested whether telomerase presence affects the F-22G3 structure. To this end, we imaged F-22G3 in the presence of catalytically active telomerase, but in the absence of deoxynucleotide triphosphates (dNTPs). Approximately 65% of F-22G3 molecules showed an abrupt drop in FRET value, from 0.53 ± 0.05 to 0.3 ± 0.1, during the 160 s after telomerase was injected into the microscopic channel containing immobilized F-22G3 (Figure 1g). The remaining 35% of molecules did not show any change in FRET signal over the observed time; it is possible that the binding reaction had not proceeded to completion within this time period, or that a subpopulation of enzyme or DNA molecules are incompetent for binding. We collected 125 molecules showing a step-wise change in FRET value and plotted the data in a FRET heat map and a histogram plot as a function of time; both plots showed a drop in mean FRET value from ∼0.53 to ∼0.3 FRET over this time (Figure 1h and Supplementary Figure 1b). We interpret this to represent telomerase binding to F-22G3 and partially opening the structure, which then remained stable in its new conformation.

We confirmed this conclusion by quantitatively analyzing the FRET changes during the transitions. For all molecules that showed a change in FRET signal over time, the frequency with which molecules transitioned between states was determined using state finding algorithm vbFRET (https://sourceforge.net). Then, the transition frequencies were plotted as a function of initial and final FRET states to obtain transition density plots (TDP) (Figure 1i). In the presence of telomerase, the TDP showed a single cluster of transitions at initial FRET ∼0.5 and final FRET ∼0.3, consistent with the shift in mean FRET in the heat map.

To examine changes in F-22G3 structure during its extension by telomerase, we performed smFRET experiments in the presence of both telomerase and dNTPs. Under these conditions, ∼65% of molecules showed a two-step drop in FRET values, from 0.53 ± 0.05 to 0.3 ± 0.1, and then to 0.15 ± 0.05 (Figure 1j-m, and Supplementary Figure 1c, d). The FRET decrease from high to low FRET states in these events was irreversible, supported by the presence of two off-diagonal clusters in the TDP (Figure 1l), suggesting a continuous irreversible unfolding of G4 structure. As a control, we performed smFRET experiments in the presence of dNTPs alone and observed no change in FRET signal (Supplementary Figure 1e). These data suggest that telomerase disrupts F-22G3 structure completely in the presence of dNTPs.

The rate of F-22G3 unfolding upon telomerase binding in the presence of dNTPs was measured by integrating the dwell time distribution at the 0.3 FRET state (τ_unfolding_; Figure 1j) and fitting the distribution to a single-exponential decay (Figure 1n). F-22G3 molecules exhibited telomerase-mediated unfolding with a rate constant of *k*_unfolding_ = 0.050 ±0.002 s^-1^ (mean ± SEM, n = 76 molecules), indicating that there is only one rate-limiting step during the unfolding process. This unfolding rate is comparable with the rate of unfolding of parallel G4 by Pif1 helicase (0.11 s^-1^) (45).

### Human telomerase also binds, unfolds and extends intermolecular parallel G-quadruplexes

We have previously demonstrated that a tetrameric, parallel G4 (46) formed from four copies of the 7-mer telomeric sequence TTAGGGT in K^+^ is highly stable but can be extended by human telomerase (34). This sequence provides the advantage that the conformation of its G4 topology is unambiguous; the sequence is too short to form intramolecular G4 structures, and can only exist as a parallel tetramer. To examine whether this tetramer is extended by telomerase using the same mechanism as an intramolecular G4, we prepared a version of this quadruplex labeled with a pair of FRET dyes and a biotin with which to immobilize the DNA. Four different strands, each consisting of the sequence TTAGGGT (here called 7GGT) and a hexa-ethylene glycol spacer, were annealed in an equimolar mixture in KCl solution. Three of these strands were 5’ modified with either AlexaFluor 555^TM^ (donor dye), AlexaFluor 647^TM^ (acceptor dye) or biotin, and the remaining strand was unmodified (Figure 2a). We assembled an equimolar mixture of each of these modified oligonucleotides to produce G4s with different combinations of modifications. The resulting mixture of G4s were all parallel-stranded, as confirmed using CD spectroscopy (Supplementary Figure 2a), and had an average melting temperature similar to that of the unmodified G4 ([7GGT]_4_; Supplementary Figure 2b). Direct telomerase activity assays demonstrated that human telomerase can extend the FRET-modified G4 construct (which we refer to as F-[7GGT]_4_) as efficiently as the unmodified [7GGT]_4_ (Figure 2b). In single-molecule microscopy analyses, we analyzed only G4 structures containing a single copy of each of the four strands, through selection during post-image processing and data analysis. Note that there are two possible orientations of the positions of the two dyes (on adjacent strands or on diagonally opposite strands, as depicted in Figure 2a), but it is unlikely that the distance between the dyes in these two conformations is sufficiently different to resolve by FRET.

**Figure 2:**
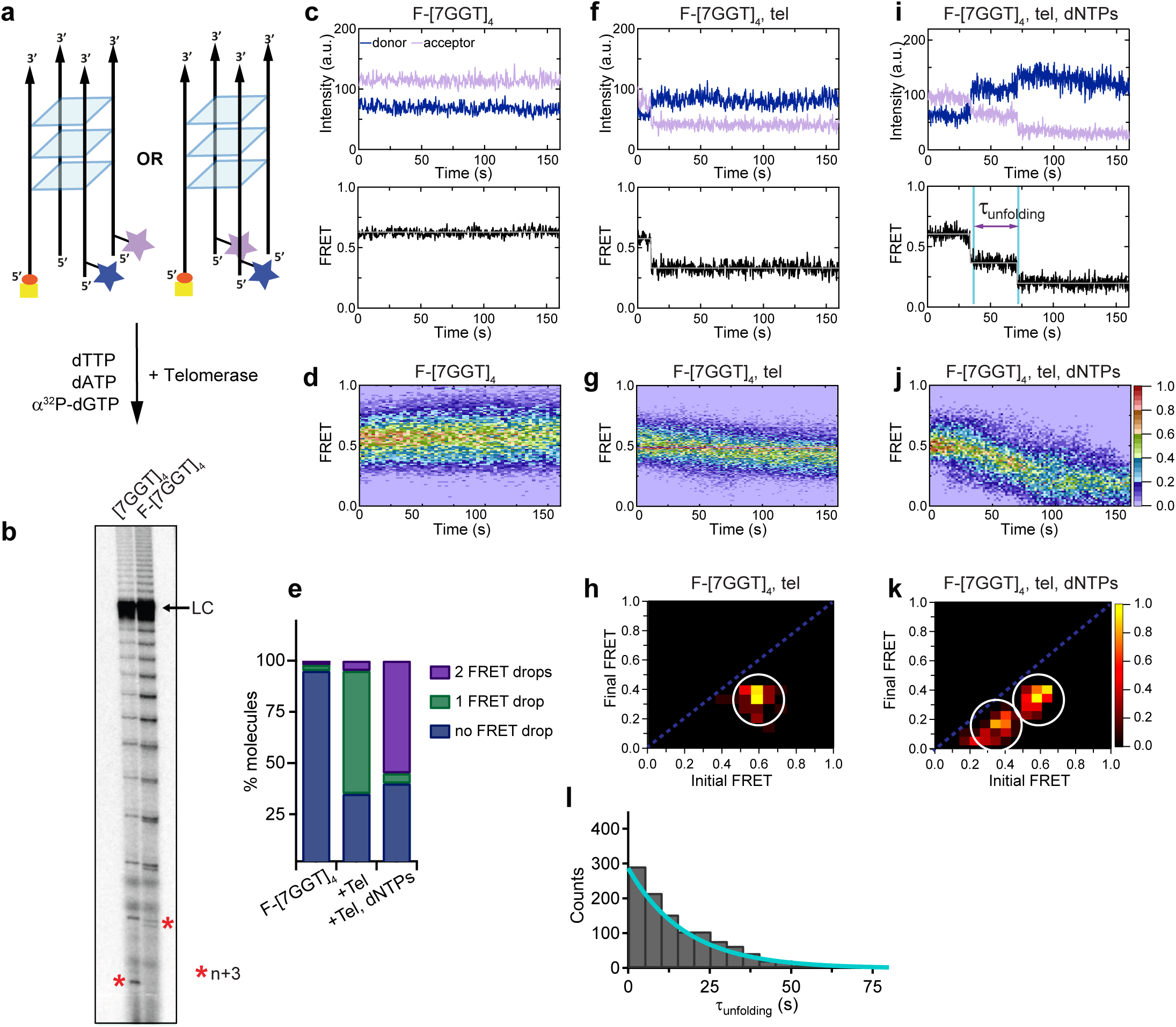
Telomerase unfolds and extends a tetrameric parallel G-quadruplex. (**a**) Schematic representation of F-[7GGT]_4_ used in smFRET studies. Blue star: AlexaFluor 555^TM^ (donor dye); purple star: AlexaFluor 647^TM^ (acceptor dye); red circle: biotin; yellow square: Neutravidin. (**b**) Telomerase extension assays using either 1 µM of unmodified [7GGT]_4_ or the version containing the FRET pair dyes (F-[7GGT]_4_); radiolabeled extension products were electrophoresed on a denaturing polyacrylamide gel (* indicates the position of n+3 products in the gel). LC: 5’-^32^P-labeled synthetic 100 nt DNA used as a recovery/loading control. (**c**),(**f**),(**i**) Representative single-molecule acceptor (purple) and donor (blue) intensities of F-[7GGT]_4_ molecules over time (top panels), and the corresponding FRET traces (bottom panels). (**d**),(**g**),(**j**) Heat maps of the distribution of FRET intensities over 0 - 150 s. For color key, see panel (**j**). Panels (**c**) and (**d**) represent F-[7GGT]_4_ alone (n = 105 molecules), (**f**) and (**g**) show F-[7GGT]_4_ in the presence of telomerase (n = 87), and (**i**) and (**j**) show F-[7GGT]_4_ in the presence of telomerase and dNTPs (n = 81). All plots include molecules collected in 4 – 6 independent experiments. (**e**) Plot of the percentage of molecules showing no change in FRET, a single FRET drop, or a two-step FRET drop, in the experiments shown above. (**h**),(**k**) Transition density plots (TDPs) depicting initial and final FRET states of all molecules that showed a change in FRET value, in the presence of telomerase (n = 65) (**h**) or telomerase and dNTPs (n = 75) (**k**). For color key, see panel (**k**). (**l**) Plot of the dwell time distribution at the 0.36 FRET state for 55 molecules from experiment in (**i-k**), representing the rate of telomerase-mediated G4 unfolding.

smFRET experiments showed that F-[7GGT]_4_ exhibits a steady FRET signal at a ratio of ∼0.6, and the FRET signal did not change over time (Figure 2c,d, Supplementary Figure 2c), indicating formation of a stable parallel intermolecular G4. Telomerase alone was sufficient for partial unwinding of parallel F-[7GGT]_4_, as demonstrated by a drop in FRET state from ∼0.6 to ∼0.4 with time (Figure 2f-h, Supplementary Figure 2d). About 60% of molecules showed this one-step drop in FRET signal upon injection of telomerase (Figure 2e). Note that the distribution of initial FRET values of those molecules that show a change in FRET value is not as broad as the FRET distribution of the whole population of molecules (compare the spread at 0 time in Figure 2g to that in Figure 2d), supporting the interpretation that those G-quadruplexes that do not bind to telomerase have folded improperly or are folding intermediates, and thus show a high level of dynamic behaviour that is too fast to resolve and results in a broadening of the FRET peak. In the case of the 60% of molecules that do show a drop in FRET signal, we interpret this to demonstrate that telomerase results in a partial separation of the FRET dyes upon binding to one strand of the tetrameric G4.

In the presence of telomerase and dNTPs, most F-[7GGT]_4_ molecules experienced a two-step drop in FRET, from 0.57 ± 0.05 to 0.36 ± 0.04, and then to 0.19 ± 0.05 (Figure 2i-k, Supplementary Figure 2e-f); this transition did not occur in the presence of dNTPs alone (Supplementary Figure 2g). The two FRET transition clusters observed by TDP analysis (Figure 2k) suggest that telomerase unfolds parallel intermolecular G4 irreversibly. The rate of unfolding of F-[7GGT]_4_ by telomerase in the presence of dNTPs was measured by plotting the dwell times of the intermediate transition states (FRET value of 0.4) and fitting them to a single exponential equation that yielded *k*_unfolding_ = 0.055 ± 0.003 s^-1^ (mean ± SEM, n = 55 molecules; Figure 2l). Thus, the rate of unfolding of [7GGT]_4_ by telomerase in the presence of dNTPs is comparable to that of F-22G3.

### Complete G-quadruplex unwinding by telomerase requires its catalytic activity

Next, we asked if either of the two step-wise drops in FRET values are dependent upon the nucleotide incorporation activity of telomerase. Synthesis activity requires three conserved aspartate residues in the reverse transcriptase domain of the TERT protein (47–50). Mutation of any one of these aspartates results in loss of telomerase catalytic activity but retention of its ability to bind to a DNA primer (51). We introduced an aspartate-to-alanine mutation at hTERT amino acid 712 and confirmed that this mutant telomerase (D712A) lost all primer extension activity (Supplementary Figure 3a). smFRET experiments with F-[7GGT]_4_ demonstrated an initial drop in FRET after addition of D712A telomerase, but no further drop in FRET was observed upon addition of dNTPs (Figure 3a-e). This result suggests that binding of telomerase to F-[7GGT]_4_, resulting in partial G4 unwinding, is independent of telomerase catalytic activity, but full unwinding of the G4 requires its extension by telomerase.

**Figure 3:**
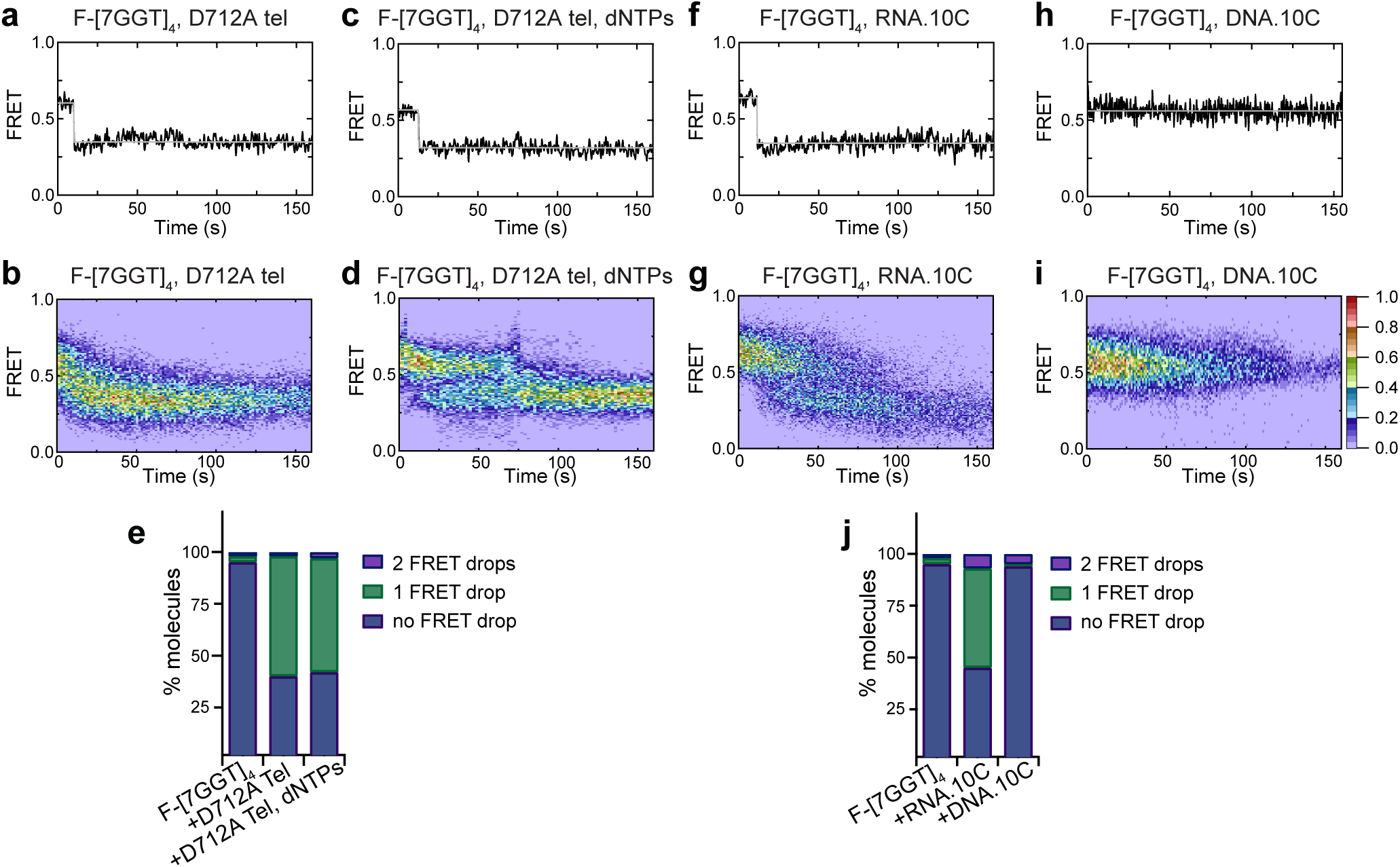
Partial unfolding of G4 does not require telomerase catalytic activity, and can be induced by the RNA template. (**a**),(**c**),(**f**),(**h**) Examples of individual F-[7GGT]_4_ FRET traces under the indicated experimental conditions over 160 s. (**b**),(**d**),(**g**),(**i**) Heat maps of the distribution of FRET trajectories over 0 - 150 s using 80, 75, 95 and 129 molecules, respectively, under the indicated experimental conditions. All plots include molecules collected in 4 – 6 independent experiments. For color key, see panel (**i**). (**e**),(**j**) Plots of the percentage of molecules showing no change in FRET, a single FRET drop, or a two-step FRET drop, in the experiments shown above.

### The RNA template sequence is involved in partial unfolding of G-quadruplex structure

It is known that the RNA template of telomerase plays an important role in recognizing and binding the telomeric end by canonical base paring. We have previously demonstrated that [7GGT]_4_ is extended by the nucleotides that would be predicted from canonical base-pairing with the RNA template (34). Therefore, we hypothesized that the RNA template binds to the G4, facilitating the opening of the structure. To test this, we performed smFRET experiments in the presence of a 10-nt RNA oligonucleotide mimicking the human telomerase template sequence, RNA.10C (Supplementary Table 1). At a high concentration of RNA.10C (500 μM), a majority of F-[7GGT]_4_ molecules executed a single-step drop in FRET from 0.57 to 0.36 (Figure 3f, g, j); thus, a short oligonucleotide resembling the telomerase template induces G4 unfolding in a similar manner to the whole telomerase enzyme. The fraction of the G4 population that showed a drop in FRET signal was dependent on the concentration of RNA.10C (Supplementary Figure 3b, c), providing evidence for direct unwinding of the G4 by the oligonucleotide, rather than passive trapping of a spontaneously unfolded structure. Supporting this conclusion, the effect on G4 structure was very specific to RNA; the presence of 500 µM of a 10-nt DNA oligonucleotide of identical sequence (DNA.10C) did not stimulate any change in the FRET signal of F-[7GGT]_4_ over the same time period (Figure 3h-j). These data suggest that the hTR template is specifically involved in binding to the G4, leading to partial opening of the structure.

### Telomerase translocation leads to complete unfolding of G-quadruplex structure

The second step of telomerase-mediated unfolding of G4 DNA requires telomerase catalytic activity (Figure 3c-e). To probe the mechanism for this, we incubated telomerase with F-22G3 in the presence of subsets of dNTPs. The first three nucleotides that are incorporated by telomerase at the 3’ end of F-22G3 are dTTP, dATP and dGTP, as dictated by the telomerase RNA template sequence (see Figure 4a, e, i). We first carried out extension reactions in the presence of only ddTTP, a chain terminator that inhibits further elongation of the 3’ end after its incorporation, and no other nucleotides (Figure 4a). Under these conditions, F-22G3 exhibited a FRET drop from ∼0.5 to ∼0.3 and remained in the ∼0.3 FRET state over the remainder of the observation time window (Figure 4b–d and Supplementary Figure 4a). A similar change in FRET from ∼0.5 to ∼0.3 was observed when the only nucleotides in the reaction were dTTP and ddATP (Figure 4f–h and Supplementary Figure 4a). However, in the presence of dTTP, dATP and ddGTP, a second step-wise drop in FRET was exhibited by F-22G3, from ∼0.3 to ∼0.15 (Figure 4j–l and Supplementary Figure 4a). These data demonstrate that complete G4 unfolding occurs after the addition of three nucleotides complementary to the template; at this point, the template boundary is reached and translocation of the DNA to the 3’ region of the template would occur (Figure 4i).

**Figure 4:**
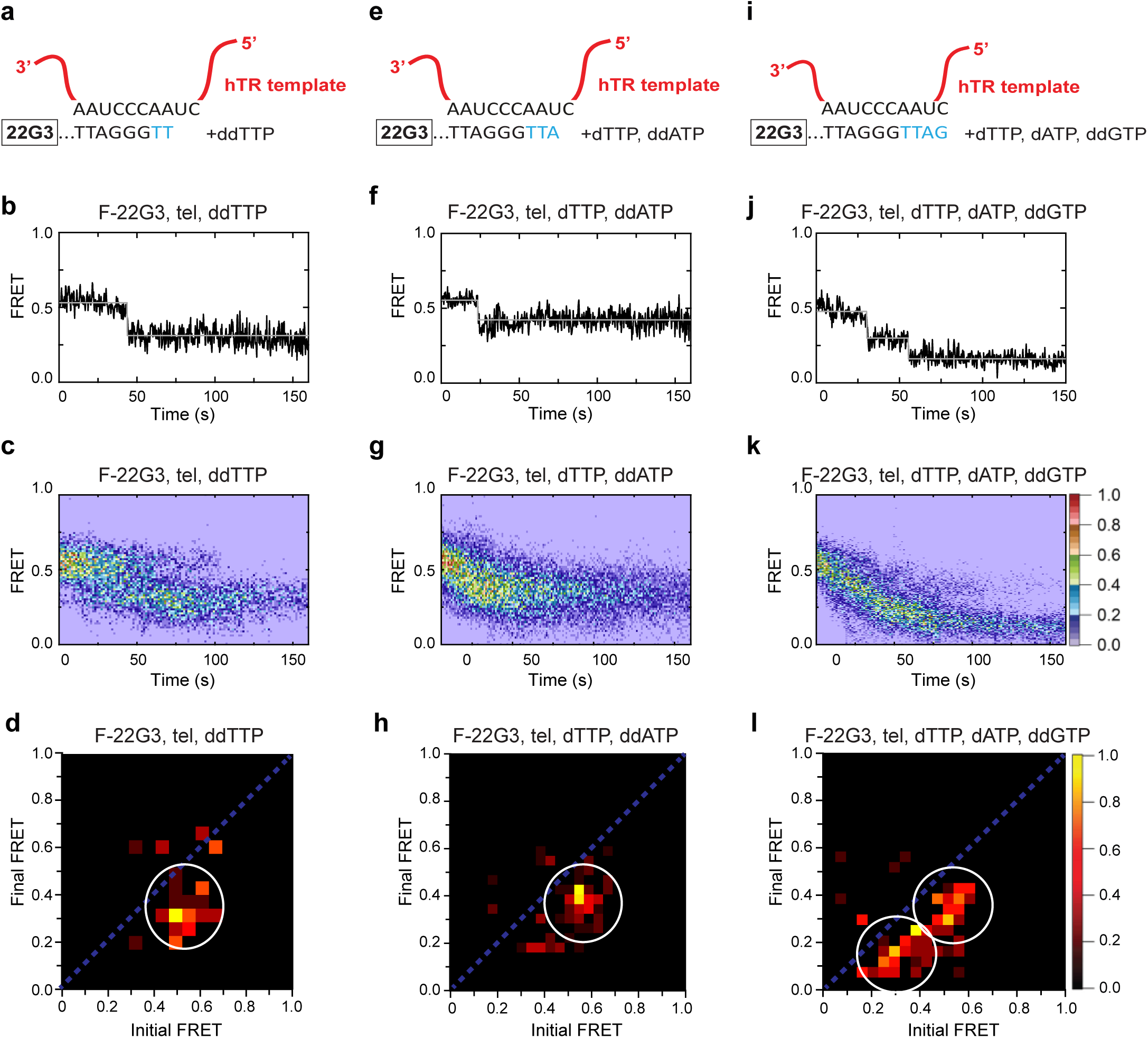
Telomerase translocation leads to complete G4 unfolding. (**a**),(**e**),(**i**) Schematic diagrams showing alignment of the telomerase template RNA with 22G3 DNA and template-directed incorporation of ddTTP (**a**), dTTP followed by ddATP (**e**) or dTTP, dATP and ddGTP (**i**). (**b**),(**f**),(**j**) Examples of individual F-22G3 FRET trajectories in the presence of telomerase and the indicated combinations of nucleotides. (**c**),(**g**),(**k**) Heat maps of the distribution of FRET intensities over 0 - 150 s in 80, 90 and 82 molecules, respectively, in the presence of telomerase and the indicated combinations of nucleotides. All plots include molecules collected in 4 – 6 independent experiments. For color key, see panel (**k**). (**d**),(**h**),(**l**) TDPs showing the changes between the initial and final FRET values of F-22G3 in the presence of telomerase and the indicated combinations of nucleotides (n = 80, 90 and 82 molecules, respectively). For color key, see panel (**l**).

### Stable intermolecular, parallel G4 is unfolded and extended by telomerase using a similar mechanism

We next asked whether telomerase-mediated unfolding of intermolecular G4 occurs via a similar mechanism to that of intramolecular G4. To address this, we performed smFRET experiments in the presence of subsets of dNTPs and observed the change in FRET signal displayed by F-[7GGT]_4_ over time (Supplementary Figure 4b-k). The FRET signal dropped in a single step, from ∼0.6 to ∼0.4, in the presence of telomerase and either ddTTP alone (Supplementary Figure 4c-e) or dTTP and ddATP (Supplementary Figure 4f-h). However, in the presence of telomerase and dTTP, dATP and ddGTP, a two-step decrease was observed, from a high FRET state (∼0.6) to ∼0.4 and then to the lowest FRET state (∼0.2) (Supplementary Figure 4b, i-k). Overall, these data demonstrate that telomerase binds and unfolds parallel G4 using a similar mechanism, whether the strand topology is intramolecular or intermolecular.

### Ligand stabilization of intramolecular parallel G4 partially inhibits but does not prevent G4 unwinding by telomerase

Antiparallel or hybrid telomeric G4 are not efficiently used as substrates by telomerase, and their stabilization with small molecule G4-binding ligands can further decrease the ability of telomerase to extend them (28, 29). We therefore sought to determine whether a ligand-mediated increase in stability of parallel G4 affects their extension by telomerase. To this end, we used three different G4-stabilizing compounds: the porphyrin N-methyl mesoporphyrin IX (NMM) (52, 53), the berberine derivative SST16 (54, 55) and the bisquinolinium compound PhenDC3 (56, 57) (Supplementary Figure 5a-c). CD spectroscopy confirmed that none of the ligands substantially changed the overall parallel G4 conformation of F-22G3 (Supplementary Figure 5d-f), and melting assays showed dramatic thermal stabilization of this G4 upon binding by all three ligands (Δ*T*_m_ of >25°C (NMM), +11°C (SST16) and >25°C (PhenDC3), under the conditions detailed in Supplementary Figure 5d-i). SST16 caused a decrease in signal of the peak at 260 nm that may be attributable to the association between the ligand and the G-quartets, resulting in slight changes in stacking interactions without a change in overall topology (58, 59). The ligands did not change the steady smFRET signal of F-22G3 in the absence of telomerase (Supplementary Figure 5j-o). In the presence of telomerase and dNTPs, the number of molecules experiencing a two-step FRET decrease was reduced by about 2-fold; nevertheless, 25 – 30% of molecules showed the same FRET decrease from ∼0.5 to ∼0.3 and then to ∼0.15 as in the absence of ligands (Figure 5a–j). The rates of telomerase-mediated unfolding of NMM-, SST16- and PhenDC3-stabilized F-22G3 were 0.018 ± 0.005 s^-1^, 0.032 ± 0.005 s^-1^, and 0.034 ± 0.004 s^-1^, respectively (mean ± SEM; Figure 5k), which were all significantly slower (*p* < 0.0001, *p* = 0.0002 and *p* = 0.0002, respectively; Student’s t-test) than in the absence of ligands (0.050 ± 0.002 s^-1^). The reduced rate of G4 unfolding provides evidence that most or all of these molecules were bound by ligands, but that ligand presence slowed the rate of their unfolding by telomerase. Consistent with these data, all three ligands partially inhibited telomerase extension of F-22G3 when incubated at the same concentrations in ensemble telomerase activity assays (Figure 5l). We therefore conclude that telomerase unfolding and extension of an intramolecular parallel G4 are partially inhibited by ligand stabilization; nevertheless, telomerase is able to overcome this stabilization and unwind a substantial proportion (25 – 30%) of molecules (Figure 5j).

**Figure 5:**
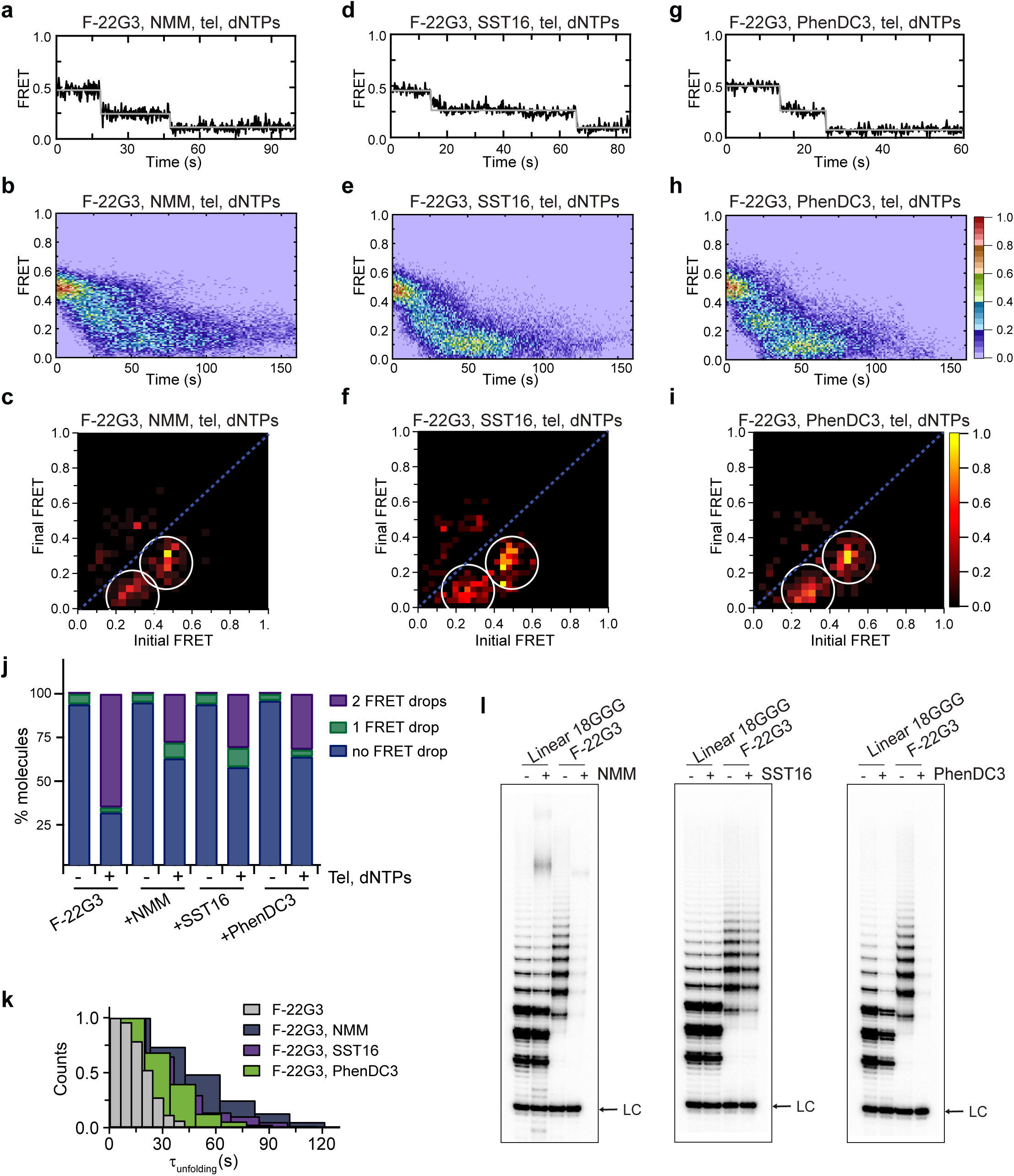
Partial inhibition of unfolding of F-22G3 by ligands NMM, SST16 and PhenDC3. (**a – i**) Representative FRET trajectories, heat maps and transition density plots of F-22G3 in the presence of NMM (**a – c**; 800 μM in folding reaction), SST16 (**d – f**; 5 μM) or PhenDC3 (**g – i**; 1 μM), in the presence of telomerase and dNTPs; n = 84 (**b**,**c**), 60 (**e**,**f**), and 77 (**h**,**i**) molecules, collected in 2-4 independent experiments. (**j**) Plot of the percentage of F-22G3 molecules showing no change in FRET, a single FRET drop, or a two-step FRET drop, when incubated with telomerase, dNTPs and the indicated ligands at the concentrations shown in (**a** – **i**). (**k**) The unfolding rate of F-22G3 in the presence of the above concentrations of NMM, SST16, PhenDC3 or no ligand, calculated by fitting the dwell time distributions of the intermediate FRET state to a single exponential equation. n = 76 (no ligand), 84 (NMM), 60 (SST16) and 77 (PhenDC3). (**l**) Telomerase extension assays using 250 nM of F-22G3 or linear Bio-L-18GGG control, in the presence or absence of NMM, SST16 or PhenDC3 at the concentrations above. For the reactions with NMM, linear DNA strands (10 μM) were incubated with 800 μM NMM prior to G4 folding, and the G4-ligand complex diluted 20-fold for the activity assay. LC: 5’-^32^P-labeled synthetic 30 nt DNA used as a recovery/loading control.

### Ligand stabilization of intermolecular parallel G4 does not inhibit telomerase activity or telomerase-mediated G4 unfolding

To determine the generality of the ability of telomerase to extend G4 in the presence of stabilizing ligands, we also incubated [7GGT]_4_ with the same three G4 ligands. CD spectroscopy confirmed that none of the ligands substantially changed the parallel G4 conformation of [7GGT]_4_ (Supplementary Figure 6a, c, e). Melting assays showed dramatic thermal stabilization of [7GGT]_4_ upon binding by all three ligands (Δ*T*_m_ of +13°C (NMM), +21°C (SST16) and +26°C (PhenDC3), under the conditions detailed in Supplementary Figure 6a-f). Telomerase activity assays were carried out at the same concentrations of [7GGT]_4_ and each ligand as used in CD analyses; surprisingly, stabilization of the G-quadruplex did not inhibit its extension by telomerase, and activity was even slightly increased when [7GGT]_4_ was stabilized by NMM (Figure 6a–c). As previously described (29), PhenDC3 caused a decrease in enzyme processivity after the addition of 4 telomeric repeats to either a linear or a G4 substrate, most likely resulting from G-quadruplex stabilization within the product DNA (Figure 6c). Concentrations of PhenDC3 higher than 1 µM caused dramatic inhibition of activity (Supplementary Fig. 6i), but again this effect was observed with both linear and G4 substrates, indicating G4-independent direct inhibition of telomerase by PhenDC3, as previously described (29). Neither NMM nor SST16 inhibited extension of linear or G4 substrates at any concentration used (Supplementary Figures 6g,h). Thus, substantial stabilization of a parallel intermolecular G4 by three different ligands did not inhibit the ability of telomerase to use the G4 as a substrate.

**Figure 6:**
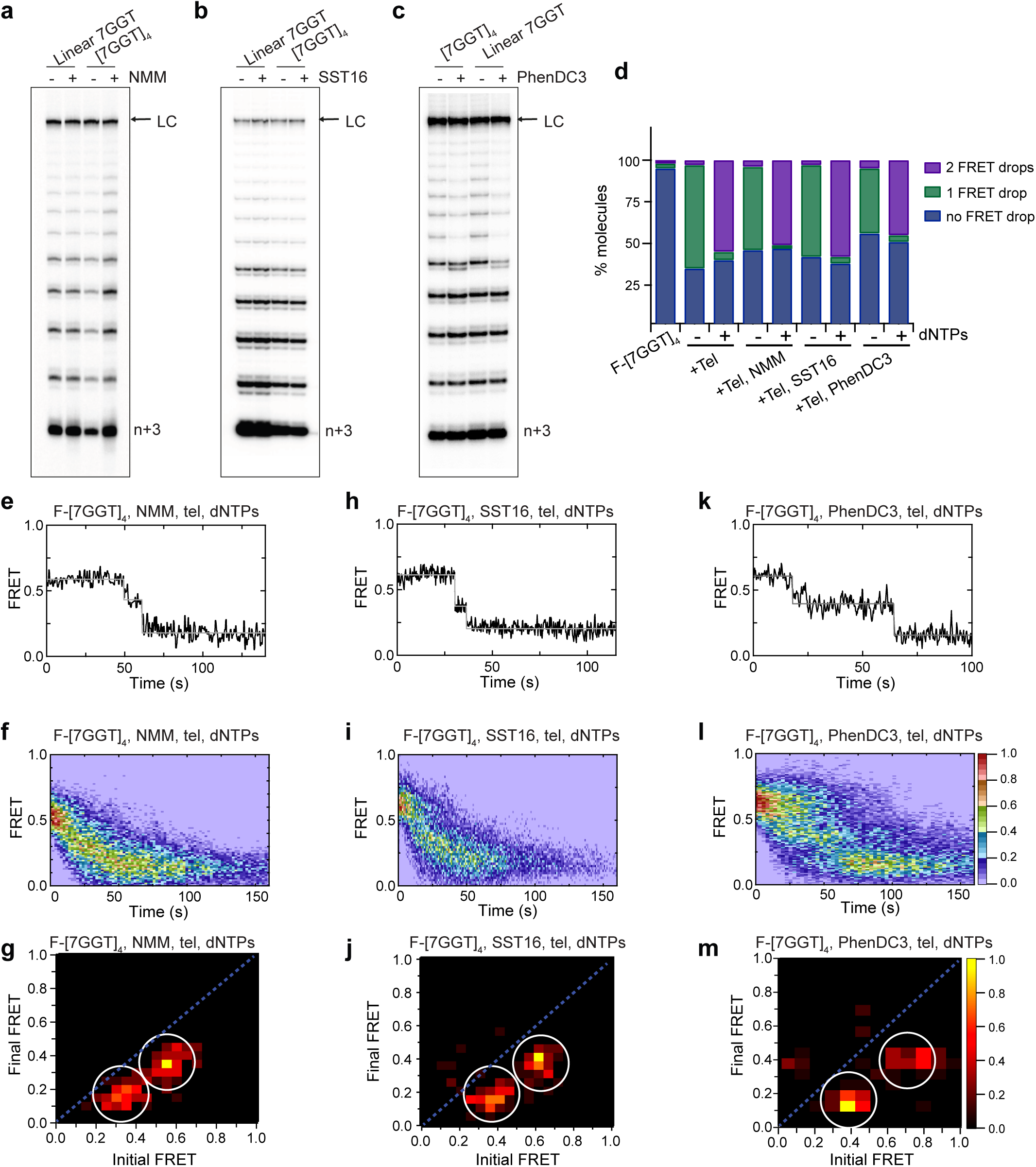
Telomerase unfolds and extends ligand-stabilized tetrameric parallel G4. (**a**),(**b**),(**c**) Telomerase extension assays using 1 µM of [7GGT]_4_ or a linear 7-mer control (or 250 nM in the reactions with PhenDC3), in the presence or absence of 40 µM NMM (**a**), 100 μM SST16 (**b**) or 1 μM PhenDC3 (**c**). For the reactions with NMM, linear 7GGT (1 mM) was incubated with 10 mM NMM prior to G4 folding, and the G4-ligand complex diluted 250-fold for the activity assay. n+3 indicates the position of the product with the first 3 nucleotides incorporated. LC: 5’-^32^P-labeled synthetic 100 nt DNA used as a recovery/loading control. (**d**) Plot of the percentage of F-[7GGT]_4_ molecules showing no change in FRET, a single FRET drop, or a two-step FRET drop, when incubated with telomerase and the indicated ligands at the concentrations shown in (**a**) – (**c**). (**e**),(**h**),(**k**) Representative single-molecule FRET trajectories of NMM-stabilized (**e**), SST16-stabilized (**h**) or PhenDC3-stabilized (**k**) F-[7GGT]_4_ in the presence of telomerase and dNTPs. (**f**),(**i**),(**l**) Heat maps of the distribution of FRET intensities over 0 - 150 s in the presence of telomerase, dNTPs and either NMM (n = 63) (**f**), SST16 (n = 90) (**i**) or PhenDC3 (n = 76) (**l**). All plots include molecules collected in 4 – 6 independent experiments. For color key, see panel (**l**). (**g**),(**j**),(**m**) TDPs showing the changes in FRET value of 70, 80 and 58 molecules from the experiments in (**e**) to (**l**).

To further understand this effect at the molecular level we performed smFRET experiments to visualize F-[7GGT]_4_ stabilized by NMM, SST16 or PhenDC3 in the presence of telomerase, with or without dNTPs. Ligand-stabilized F-[7GGT]_4_ exhibited a constant FRET signal at 0.57 (Supplementary Figure 6j-o), but in the presence of telomerase, a single-step FRET decrease from 0.57 to 0.36 was observed in a majority of single-molecule traces (Figure 6d, Supplementary Figures 6p-u). These data suggest that ligand binding to F-[7GGT]_4_ did not prevent telomerase from inducing a conformational change in the G4, as occurs in the absence of ligand. When F-[7GGT]_4_ was incubated with both telomerase and dNTPs in the presence of each ligand, most molecules showed a unidirectional two-step decrease in FRET value, from 0.57 to 0.36, and then to 0.19 (Figure 6d-m). We assessed the F-[7GGT]_4_ unfolding rate in the presence of the ligands by measuring the dwell time of each molecule in the transient intermediate state (0.36 FRET) and fitting time distributions with a single exponential equation. The rates of telomerase-mediated unfolding of NMM-, SST16- and PhenDC3-stabilized G4 were 0.085 ± 0.004 s^-1^, 0.088 ± 0.004 s^-1^, and 0.032 ± 0.001 s^-1^, respectively (mean ± SEM; Supplementary Figure 6v). Thus, while PhenDC3 resulted in a decrease in the unfolding rate of this G-quadruplex, in the presence of NMM and SST16 unfolding occurred significantly faster (*p* < 0.0001; Student’s t-test) than in the absence of ligands (0.055 ± 0.002 s^-1^). Together, these data demonstrate that neither unwinding nor extension of parallel intermolecular G4 by telomerase were inhibited by ligand-mediated stabilization of the G4.

## Discussion

Previously, we have shown that highly purified human telomerase can disrupt parallel intermolecular G4 structures and then extend them in a processive manner (34). Here, we provide the first mechanistic details of telomerase resolution of parallel G4 structures. Our data demonstrate that G4 unfolding occurs in three major steps: i) the telomerase template RNA actively invades the G-quartets in order to hybridize with the G-rich DNA, causing partial opening of the G4 structure; ii) telomerase extends the 3’ end of this partially unwound structure, adding single nucleotides according to the RNA template, and iii) translocation of telomerase once nucleotide addition has reached the template boundary triggers complete unfolding of the G4 structure (Figure 7). It has previously been difficult to distinguish whether it is the parallel or intermolecular nature of G4 structures that allows their recognition by telomerase. Here, we overcome this difficulty by exploiting the stabilization of intramolecular parallel G4 conferred by the modified nucleotide 2’F-araG (44), enabling the demonstration that inter- and intramolecular parallel G4 are extended by telomerase using the same 3-step mechanism. These data reveal human telomerase to be a parallel G4 resolvase.

**Figure 7:**
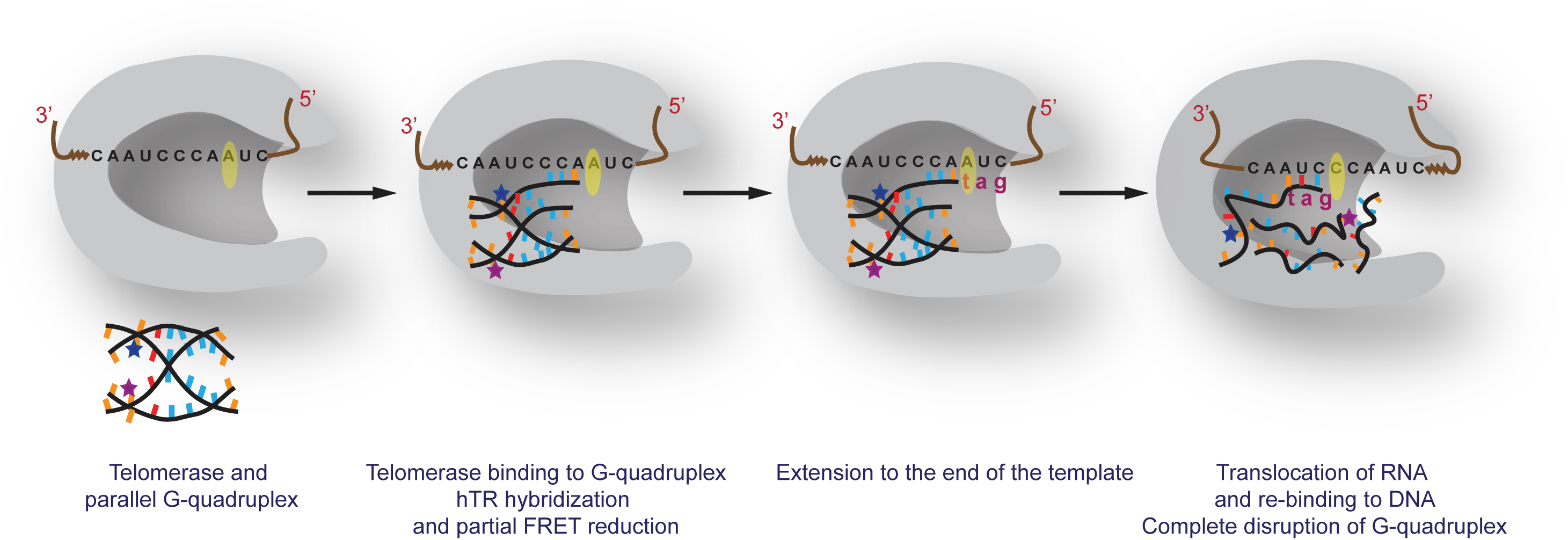
A proposed model for parallel G4 extension and unfolding by human telomerase. See text for details.

Several other proteins have been shown to assist telomerase extension of telomeric DNA by resolving antiparallel G4 in the substrate, including telomeric protein POT1 (26, 60), a splice variant of hnRNP A2 (61), and the ciliate protein StyRecQL (62, 63). In contrast, we have verified that highly purified telomerase, free of contaminating helicases or other proteins, is able to extend [7GGT]_4_ (34). Therefore, we conclude that telomerase itself has the ability to unwind parallel G4, without the assistance of interacting helicases.

Telomerase partially unfolds parallel G4, independently of telomerase catalytic activity, as indicated by the drop in FRET signal upon telomerase association with G4 (Figure 3a-e). The resulting stable intermediate FRET value is consistent with a partial separation of the FRET dyes in the absence of complete strand dissociation; we interpret these data to indicate that binding of telomerase RNA to the nucleotides at the 3’ end of one DNA strand allows RNA-DNA hybridization to occur, but the dyes are still sufficiently close together to give a non-zero FRET signal (Figure 7). To investigate whether the template RNA of telomerase plays a critical role in this initial G4 unfolding, we carried out experiments in the presence of a short RNA oligonucleotide with a sequence complementary to the telomere repeat, mimicking the template region of hTR. This RNA molecule was able to partially unfold the structure in a similar manner to the whole enzyme. This observation suggests that it is the RNA component of telomerase, rather than the hTERT protein, that promotes the initial unwinding of parallel G4, followed by hybridization to the 3’ end. A DNA oligonucleotide of identical sequence did not promote any G4 unwinding (Figure 3h-j); if oligonucleotide binding was due to passive trapping of a spontaneously-unfolding G4, this would also have been observed in the presence of a DNA C-strand. There are two lines of evidence to suggest that the difference in behaviour of the DNA and RNA oligonucleotides cannot simply be explained by the potentially higher stability of a DNA-RNA hybrid. Firstly, for this to be the case, there would need to be initial binding of either the DNA or the RNA oligonucleotide to the G4 DNA in order to form a hybrid. However, the tetrameric parallel G4s used in these experiments is highly stable; it did not demonstrate any spontaneous unwinding during the observation time of the smFRET experiments, and in our previous work we have shown that the parallel tetrameric G4 did not show any unwinding over the time scale of hours (34). Secondly, the concentration-dependent unfolding by the RNA oligonucleotide (Supplementary Figure 3b, c) also supports an active invasion mechanism. A similar template-mimicking RNA was unable to form a DNA-RNA hybrid with an antiparallel G4 of similar stability to the parallel G4 used in the present study (64), demonstrating that this is a specific interaction between this RNA and parallel G4 structures. Single-molecule FRET studies have illuminated the mechanisms used by other G4-unwinding proteins; for example, the helicase RHAU/DHX36 was shown to stack on the top of the G4 plane and bind to the exposed 3’ single stranded DNA tail (65), whereas the replication protein RPA has been proposed to bind to G4 structures via their exposed loops (66). Our data demonstrate that telomerase possesses a unique mechanism of G4 resolution, mediated by its integral RNA subunit.

Telomerase extends the 3’ end of parallel G4 while the G4 remains partially structured, as demonstrated by the intermediate FRET state remaining unchanged until the hTR template boundary was reached. Upon translocation, complete G4 unfolding occurs, represented by a second step-wise drop in FRET signal. One possible explanation for this observation is that the G4 structure needs to pass through the telomerase DNA-binding cleft during the conformation changes required for telomerase translocation, and the resulting steric hindrance results in complete dissociation of the parallel telomeric G4 (Figure 7). It has also been proposed that dissociation of the RNA-DNA hybrid occurs outside the active site, and the realigned hybrid is then re-captured into the active site (67); it is plausible that this recapturing leads to G4 dissociation. Our smFRET experiments do not address where RNA/DNA association and dissociation occur, but we hypothesize that steric hindrance encountered by the G4 structure may play a major role in the complete unfolding of the G4, whether this occurs within the catalytic active site or upon re-entry.

We used the G4-stabilizing ligands NMM, SST16 and PhenDC3, all of which bind to parallel G4 (53, 54, 68), to understand whether telomerase extension of these structures can be modulated. Despite substantial thermal stabilization of the G4, these ligands did not inhibit telomerase extension of intermolecular parallel telomeric G4, with two of them instead slightly accelerating G4 unfolding. The ligands did reduce the number of intramolecular parallel G4 molecules unwound by telomerase; nevertheless, some molecules were unwound by telomerase despite ligand binding, using the same two-step mechanism as in the absence of ligands. The fact that telomerase was able to extend parallel G4 even after substantial thermal stabilization of the structure using either small molecule ligands (Figures 5, 6) or chemical modification (Figure 1) provides further evidence that telomerase is not simply exploiting inherent instability in the G4 structure to passively extend spontaneously unwound DNA, but is instead actively involved in G4 unwinding.

NMM has been demonstrated to bind to parallel telomeric G4 by stacking on the outermost G-quartets (69), which would be predicted to interfere with telomerase extension. This observation suggests that, like other proteins such as RHAU helicase (30, 70), telomerase is able to dislodge ligands from the tetramolecular G4 structure while unfolding it. This displacement activity is structure dependent, since telomerase is less able to dislodge ligands from a parallel intramolecular G4, possibly due to the presence of DNA loops. Such a displacement mechanism would have implications for interpreting the biological effects of G4-stabilizing ligands in cell-based studies, since any effects on telomerase action and telomere length will depend on the conformational specificity of the ligand.

There is growing evidence for G4 formation at telomeres (8, 34, 71–74). In human cells, it is not known if they form at the 3’ end of the telomeric overhang, internally, or a combination of both, or which conformation of G4 occurs at these locations *in vivo*. While in-cell NMR studies of transfected telomeric oligonucleotides detected hybrid and antiparallel structures (75), a study using parallel-specific antibodies reported detection of parallel telomeric G-quadruplexes in human cells (76) and a parallel-specific ligand induces G4 formation at human telomeres (34). It is also possible that different conformations exist at different telomeric overhangs within the same cell. Intermolecular parallel G4 may also form between telomeric overhangs; it has been suggested that tetrameric parallel G-quadruplexes could be responsible for the correct alignment of four chromatids during meiosis (77). Regardless of this uncertainty, our data unequivocally demonstrate that human telomerase interacts directly with parallel G4 structures and resolves them. The conserved ability of human and ciliate telomerase enzymes to unwind these structures suggests that parallel G-quadruplexes form at telomeric overhangs *in vivo*, and do not form a barrier to telomerase extension.

## Methods

### Oligonucleotides and G-quadruplex preparation

Most DNA oligonucleotides (Supplementary Table 1) were purchased from Integrated DNA Technologies and RNA oligonucleotides from Dharmacon, with purification by high performance liquid chromatography (HPLC). Oligonucleotides 22G0, 22G3, and 22G0+tail were synthesized as previously described (44). Synthesis of (C5-alkylamino)-dT-41G3 (the equivalent of 22G3+tail prior to AlexaFluor 555^TM^ conjugation) was performed on an ABI 3400 DNA synthesizer (Applied Biosystems) at 1 μmol scale on Unylinker CPG solid support (Chemgenes). Conjugation of (C5-alkylamino)-dT-41G3 to AlexaFluor 555^TM^ NHS Ester to produce the desired 22G3+tail oligonucleotide was achieved following the standard protocol by Sigma-Aldrich (Protocol for Conjugating NHS-Ester Modifications to Amino-Labeled Oligonucleotides). Briefly, a solution of (C5-alkylamino)-dT-41G3 (200 μL, 0.3 mM) in sodium tetraborate decahydrate buffer (0.091 M, pH 8.5) was combined with a solution of AlexaFluor 555 in anhydrous DMSO (50 μL, 8 mM) and the reaction mixture was left shaking at room temperature for 2.5 hours. Samples were evaporated to dryness and the 22G3+tail product was purified by anion exchange HPLC as described (44). The peak of (C5-alkylamino)-dT-41G3 eluted between 24 and 26 min, while the desired product peak of 22G3+tail eluted between 27 and 28.5 min. Based on the area of the two peaks, the yield of the conjugation reaction is approximately 25%. The collected sample of 22G3+tail was desalted on NAP-25 desalting columns according to the manufacturer’s protocol. The maximum absorbance of AlexaFluor 555 (555 nm) in the purified 22G3+tail was verified with UV-Vis spectroscopy. The mass of 22G3+tail was verified by high resolution liquid chromatography– mass spectrometry (LC–MS; 14,000.23 m/z).

Intramolecular G4 formation using 22G0 and derivatives (Supplementary Table 1) was performed at a DNA concentration of 10 μM in 20 mM potassium phosphate, 70 mM potassium chloride pH 7, 1 mM MgCl_2_, by heat denaturing for 10 min at 90 °C, allowing the DNA to cool slowly (∼1 h) to 25 °C and equilibration at this temperature for 12-16 h. Formation of fluorescently-labeled F-22G3 with a duplex tail (Figure 1c) involved incubation of 10 µM each of oligonucleotides 22G3+tail and 647-Strand2 (Supplementary Table 1) under the same folding conditions as given for 22G0.

7GGT and its labeled derivates were combined at a final concentration of 1 mM in K^+^ hTel buffer (50 mM Tris-HCl, pH 8, 1 mM MgCl_2_, 150 mM KCl) and heat denatured for 5 min at 95 °C. They were allowed to cool slowly (∼1 h) to 25 °C and left to equilibrate at this temperature for 72 h. Intermolecular G4 formation was confirmed by native gel electrophoresis followed by staining with SYBR® Gold (Life Technologies), as described (34). DNA concentrations were determined by UV absorbance at 260 nm (extinction coefficients in Supplementary Table 1). Concentrations of G-quadruplexes are given as the concentration of assembled complexes (i.e. taking strand stoichiometry into account). Folded G-quadruplexes were stored at 4 °C until use.

### Circular dichroism

Circular dichroism (CD) spectra were recorded at 25 °C on either an Aviv 215S or a JASCO J-810 CD spectrometer equipped with Peltier temperature controllers. G-quadruplex samples of the desired conformation were prepared at 250 nM - 20 µM in their folding buffers. Three to four scans were accumulated over the indicated wavelength ranges in a 0.1 cm or 1 cm path length cell. Parameters used with the Aviv CD spectrometer included a time constant of 100 ms, averaging time 1 s, sampling every 1 nm, and bandwidth 1 nm, while the JASCO CD spectrometer was used with a scan rate of 100 nm/min and a response time of 2.0 s. Buffers alone were also scanned and these spectra subtracted from the average scans for each sample. CD spectra were collected in units of millidegrees, normalized to the total species concentrations and expressed as molar ellipticity units (deg × cm^2^ dmol^-1^). Data were smoothed using the Savitzky-Golay function within the JASCO graphing software, or the smoothing function within GraphPad Prism. For thermal stability analysis, the samples were scanned using the above parameters, but with a fixed wavelength (260 nm) over increasing temperature (25 °C to 100 °C), at a rate of ∼1 °C/min. For reactions including SST16 or PhenDC3 (prepared as described (54, 56)), the ligand was incubated with the folded DNA substrate at 25 °C for 30 min prior to CD spectroscopy. For reactions including NMM (Frontier Scientific, USA), the ligand was incubated with the DNA prior to G-quadruplex folding and the G4-ligand complex diluted prior to CD spectroscopy. Concentrations of ligands and G4 DNA used for each experiment are given in the figure legends.

### Preparation of telomerase

Human telomerase was overexpressed in HEK293T cells and purified as described (34, 78). Briefly, plasmids encoding hTERT, hTR and dyskerin (available from the authors with an accompanying Materials Transfer Agreement) were transiently transfected into HEK293T cells growing in 20 L bioreactors using polyethylenimine and cells harvested 4 days later. Cell lysates were clarified, ribonucleoprotein complexes enriched with MgCl_2_, and telomerase immunoprecipitated with a sheep polyclonal hTERT antibody, raised against hTERT amino acids 276-294 (ARPAEEATSLEGALSGTRH) (79); available from Abbexa Ltd., Cambridge, UK (catalogue number abx120550). Telomerase was eluted by competitive elution with the same peptide in 20 mM HEPES-KOH (pH 8), 300 mM KCl, 2 mM MgCl_2_, 0.1% v/v Triton X-100 and 10% v/v glycerol. Fractions were assayed for telomerase concentration by dot-blot northern against hTR (78), and equal amounts of enzyme (∼1.5 nM) used in each activity assay.

### Telomerase activity assays

The following reaction was prepared to give 20 µL per sample: 250 nM - 2 µM of the specified oligonucleotide (concentrations given in figure legends), 20 mM HEPES-KOH (pH 8), 2 mM MgCl_2_, 150 mM KCl, 5 mM dithiothreitol, 1 mM spermidine-HCl, 0.1% v/v Triton X-100, 0.5 mM dTTP, 0.5 mM dATP, 4.6 µM nonradioactive dGTP and 0.33 µM [α-^32^P]dGTP at 20 mCi mL^-1^, 6000 Ci mmol^-1^ (PerkinElmer Life Sciences). For reactions including SST16 or PhenDC3, the ligand was incubated with the folded DNA substrate at 25 °C for 30 min prior to adding other components. For reactions including NMM, the ligand was incubated with the DNA prior to G-quadruplex folding; concentrations are given in figure legends. Telomerase activity assays were initiated by adding purified human telomerase to ∼1.5 nM, and incubating at 37 °C for 1 h. The reaction was quenched by the addition of 20 mM EDTA and 1-2 x 10^3^ cpm of a 5’-^32^P-labeled synthetic 100-mer, 30-mer or 12-mer DNA (as indicated in figure legends) as an internal recovery standard. Products of telomerase extension were recovered as described, either with phenol/chloroform extraction followed by ethanol precipitation (34), or, for biotinylated substrates, by recovery with magnetic streptavidin beads (78). The solution was heated at 90 °C for 5 min, and 3 μL was electrophoresed over a 10% polyacrylamide sequencing gel (0.2 mm thick x 40 cm length x 35 cm width, 32-well comb) run in 1 × TBE/8 M urea at 85 W. The gel was transferred to filter paper, dried for 30 min at 80 °C, exposed to a PhosphorImager screen, visualized on a Typhoon FLA9500 scanner (GE Healthcare Lifesciences) and analyzed using ImageQuant^TM^ software.

### Single-molecule Fluorescence imaging and data analysis

#### Microscope setup for FRET imaging

A home-built objective-type total internal reflection fluorescence (TIRF) microscope based on an Olympus IX-71 model was used to record single-molecule movies. A Coherent Sapphire green (532 nm) laser was used to excite donor molecules at an angle of TIRF by focusing on a 100X oil immersed objective. FRET was measured by excitation with a 532 nm laser and the emissions at 565 and 665 nm were collected using a band pass filter at 560-600 nm and a long pass filter at 650 nm. Scattered light was removed by using a 560 nm long pass filter. AlexaFluor 555 and AlexaFluor 647 signals were separated by 638 nm dichroic using photometrics dual view (DV-2) and both signals were focused onto a charge-coupled device (CCD) camera (Hamamatsu C9 100-13), simultaneously. Data were collected at 5 frames per second.

#### Sample preparation for FRET experiments

Quartz coverslips were treated with 100% ethanol and 1 mM KOH. Then, aminosilanization of coverslips was carried out in a 1% v/v (3-Aminopropyl)triethoxy silane (Alfa Aesar, A10668, UK) solution in acetone. PEGylation was carried out by incubating a mixture of biotinPEG-SVA and mPEG-SVA (Laysan Bio, AL, USA) at a ratio of 1:20 prepared in 0.1 M NaHCO_3_ solution on the top of a silanized coverslip for at least 3-4 h. Finally, PEGylated coverslips were stored under dry nitrogen gas at −20 °C.

#### Single-molecule experiments

Immuno-pure Neutravidin solution was prepared in K^+^ buffer (10 mM Tris-HCl (pH 8), 1 mM MgCl_2_ and 150 mM KCl) and spread on the top of a dry PEGylated coverslip followed by a 10 min incubation. Sample flow chambers were created by sandwiching polydimentylsiloxane (PDMS) on top of the neutravidin coated coverslip. Then, blocking buffer (K^+^ buffer with 1% Tween-20) was injected into the channel in order to reduce non-specific binding of proteins on the surface, followed by 10-15 min incubation. A 50 pM solution of biotinylated FRET G-quadruplex substrate was prepared in K^+^ buffer and 200 µL injected into the flow chamber using a syringe pump (ProSense B.V.) followed by incubation for 10 min. Unbound sample was washed off in K^+^ buffer. Movies were recorded at room temperature (20 ± 1°C) for 3-5 minutes in oxygen-scavenging system (OSS) consisting of protocatechuic acid (PCA, 2.5 mM) and protocatechuate-3,4-dioxigenase (PCD, 50 nM) to reduce photo-bleaching of the fluorophores, and 2 mM Trolox to reduce photo-blinking of dyes. For experiments in the presence of enzyme, 200 µL of telomerase (0.5 nM; expressed and purified as described above) and/or dNTPs or ddNTPs (0.5 mM) were injected into the microscopic channel containing immobilized G4 DNA while the movie was recorded continuously. The buffer reached the microscopic channel at ∼10 s and the movie was collected until all acceptor molecules photobleached. For reactions including SST16 or PhenDC3, the ligands were incubated with the folded DNA substrate at 25 °C for 30 min prior to dilution for sample injection. For reactions including NMM, the ligand was incubated with the DNA prior to G-quadruplex folding, and the G4-ligand complex diluted to 50 pM G4 prior to injection. Concentrations of DNA and ligands combined for each experiment are given in the figure legends.

#### Data analysis

Single-molecule intensity time trajectories were generated in IDL (software available at https://cplc.illinois.edu/software/) and these trajectories were analyzed in MATLAB using home-written scripts (available from the authors upon request). Approximate FRET value is measured as the ratio of acceptor intensity to the sum of the donor and acceptor intensities after correcting cross talk between donor and acceptor channels. The collected FRET traces were further analyzed using the vbFRET algorithm (https://sourceforge.net) to find possible FRET states and transition frequency among these FRET states with a Hidden Markov Model (HMM).

To measure rate constants for G4 unfolding, integrated dwell time histograms were constructed and fitted to a single exponential decay function of the form:

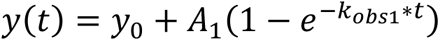

where A_1_ is the fraction of decay population and *k*_obs1_ is the apparent observed rate constant.

#### Gaussian fitting of the cumulative FRET histograms

Multiple-Gaussian fit model was applied to FRET histograms generated by binning many FRET trajectories to obtain mean FRET values.

## Acknowledgements

We thank Omesha Perera for plasmid construction, George Lovrecz and Tram Phan for production of telomerase-expressing cells, Timothy Adams for providing pAPEX vectors and for consultation on overexpression of telomerase, and Corinne Getta for preparation of PhenDC3. The work in the Bryan laboratory was supported by Cancer Council NSW project grants RG 11-07 and RG 16-10, a Denise Higgins PhD Scholarship and a Kids Cancer Alliance PhD Top Up Scholarship from Cancer Institute NSW (to A.L.M.). The work in the van Oijen laboratory was supported by an Australian Laureate Fellowship to A.M.v.O. (FL140100027). S.B.C. was supported by the Ernest and Piroska Major Foundation. M.L.B. was supported by an Australian Postgraduate Award. Funding in the Damha laboratory was provided by the National Science and Engineering Council of Canada (Discovery NSERC grant). S.S. was supported by Center of Excellence for Innovation in Chemistry (PERCH-CIC) and Research Unit of Natural Products and Organic Synthesis for Drug Discovery (NPOS 405/2560).

## Author contributions

The smFRET assay was designed and the study conceived by B.P., A.L.M., A.M.v.O. and T.M.B.; smFRET experiments were performed and analyzed by B.P. and A.M.v.O.; telomerase was purified and ensemble telomerase assays were performed and analyzed by A.L.M., C.G.T., T.M.B. and S.B.C.; modified oligonucleotide synthesis and characterization were performed by H.A.A., R.E.K., C.G. and M.J.D.; synthesis and characterization of G4 ligands was performed by S.S., K.I., M.L.B., J.L.B. and M.-P. T.-F.; the manuscript was written by B.P. and T.M.B.; and all authors edited the manuscript.

## Code availability

The software used for data analysis is freely available at https://cplc.illinois.edu/software/ (IDL) and https://sourceforge.net (vbFRET). The custom MATLAB script used for trajectory analysis is available from the authors upon request.

## References

1. Rhodes D, Lipps HJ. G-quadruplexes and their regulatory roles in biology. Nucleic Acids Res. 2015;43(18):8627–37.

2. Maizels N, Gray LT. The G4 genome. PLoS Genet. 2013;9(4):e1003468.

3. Bochman ML, Paeschke K, Zakian VA. DNA secondary structures: stability and function of G-quadruplex structures. Nat Rev Genet. 2012;13(11):770–80.

4. Huppert JL, Balasubramanian S. Prevalence of quadruplexes in the human genome. Nucleic Acids Res. 2005;33(9):2908–16.

5. S. Neidle SB. Quadruplex Nucleic Acids: Royal Society of Chemistry; 2006.

6. Williamson JR, Raghuraman MK, Cech TR. Monovalent cation-induced structure of telomeric DNA: the G-quartet model. Cell. 1989;59(5):871–80.

7. Blackburn EH. Structure and function of telomeres. Nature. 1991;350:569.

8. Biffi G, Tannahill D, McCafferty J, Balasubramanian S. Quantitative visualization of DNA G-quadruplex structures in human cells. Nat Chem. 2013;5(3):182–6.

9. Gomez D, Wenner T, Brassart B, Douarre C, O’Donohue MF, El Khoury V, … Riou JF. Telomestatin-induced telomere uncapping is modulated by POT1 through G-overhang extension in HT1080 human tumor cells. J Biol Chem. 2006;281(50):38721–9.

10. Burge S, Parkinson GN, Hazel P, Todd AK, Neidle S. Quadruplex DNA: sequence, topology and structure. Nucleic Acids Res. 2006;34(19):5402–15.

11. Renciuk D, Kejnovska I, Skolakova P, Bednarova K, Motlova J, Vorlickova M. Arrangements of human telomere DNA quadruplex in physiologically relevant K+ solutions. Nucleic Acids Res. 2009;37(19):6625–34.

12. Miller MC, Buscaglia R, Chaires JB, Lane AN, Trent JO. Hydration is a major determinant of the G-quadruplex stability and conformation of the human telomere 3’ sequence of d(AG3(TTAG3)3). Journal of the American Chemical Society. 2010;132(48):17105–7.

13. Heddi B, Phan AT. Structure of human telomeric DNA in crowded solution. Journal of the American Chemical Society. 2011;133(25):9824–33.

14. Olovnikov AM. [Principle of marginotomy in template synthesis of polynucleotides]. Dokl Akad Nauk SSSR. 1971;201(6):1496–9.

15. Watson JD. Origin of concatemeric T7 DNA. Nat New Biol. 1972;239(94):197–201.

16. Harley CB, Futcher AB, Greider CW. Telomeres shorten during ageing of human fibroblasts. Nature. 1990;345(6274):458–60.

17. Greider CW, Blackburn EH. The telomere terminal transferase of Tetrahymena is a ribonucleoprotein enzyme with two kinds of primer specificity. Cell. 1987;51(6):887–98.

18. Greider CW, Blackburn EH. Identification of a specific telomere terminal transferase activity in Tetrahymena extracts. Cell. 1985;43(2 Pt 1):405–13.

19. Greider CW, Blackburn EH. A telomeric sequence in the RNA of Tetrahymena telomerase required for telomere repeat synthesis. Nature. 1989;337(6205):331-7.

20. Greider CW. Telomerase is processive. Mol Cell Biol. 1991;11(9):4572–80.

21. Hiyama E, Hiyama K. Telomere and telomerase in stem cells. Br J Cancer. 2007;96(7):1020–4.

22. Podlevsky JD, Chen JJ. It all comes together at the ends: telomerase structure, function, and biogenesis. Mutat Res. 2012;730(1-2):3–11.

23. Wyatt HD, West SC, Beattie TL. InTERTpreting telomerase structure and function. Nucleic Acids Res. 2010;38(17):5609–22.

24. Smith JS, Chen Q, Yatsunyk LA, Nicoludis JM, Garcia MS, Kranaster R, … Johnson FB. Rudimentary G-quadruplex-based telomere capping in Saccharomyces cerevisiae. Nat Struct Mol Biol. 2011;18(4):478–85.

25. Zahler AM, Williamson JR, Cech TR, Prescott DM. Inhibition of telomerase by G-quartet DNA structures. Nature. 1991;350(6320):718–20.

26. Zaug AJ, Podell ER, Cech TR. Human POT1 disrupts telomeric G-quadruplexes allowing telomerase extension in vitro. Proc Natl Acad Sci U S A. 2005;102(31):10864–9.

27. Gomez DL, Armando RG, Cerrudo CS, Ghiringhelli PD, Gomez DE. Telomerase as a Cancer Target. Development of New Molecules. Curr Top Med Chem. 2016;16(22):2432–40.

28. Sun D, Thompson B, Cathers BE, Salazar M, Kerwin SM, Trent JO, … Hurley LH. Inhibition of human telomerase by a G-quadruplex-interactive compound. J Med Chem. 1997;40(14):2113–6.

29. De Cian A, Cristofari G, Reichenbach P, De Lemos E, Monchaud D, Teulade-Fichou MP, … Mergny JL. Reevaluation of telomerase inhibition by quadruplex ligands and their mechanisms of action. Proc Natl Acad Sci U S A. 2007;104(44):17347–52.

30. Tippana R, Hwang H, Opresko PL, Bohr VA, Myong S. Single-molecule imaging reveals a common mechanism shared by G-quadruplex-resolving helicases. Proc Natl Acad Sci U S A. 2016;113(30):8448–53.

31. Budhathoki JB, Maleki P, Roy WA, Janscak P, Yodh JG, Balci H. A Comparative Study of G-Quadruplex Unfolding and DNA Reeling Activities of Human RECQ5 Helicase. Biophys J. 2016;110(12):2585–96.

32. Liu JQ, Chen CY, Xue Y, Hao YH, Tan Z. G-quadruplex hinders translocation of BLM helicase on DNA: a real-time fluorescence spectroscopic unwinding study and comparison with duplex substrates. Journal of the American Chemical Society. 2010;132(30):10521–7.

33. London TB, Barber LJ, Mosedale G, Kelly GP, Balasubramanian S, Hickson ID, … Hiom K. FANCJ is a structure-specific DNA helicase associated with the maintenance of genomic G/C tracts. J Biol Chem. 2008;283(52):36132–9.

34. Moye AL, Porter KC, Cohen SB, Phan T, Zyner KG, Sasaki N, … Bryan TM. Telomeric G-quadruplexes are a substrate and site of localization for human telomerase. Nat Commun. 2015;6:7643.

35. Oganesian L, Moon IK, Bryan TM, Jarstfer MB. Extension of G-quadruplex DNA by ciliate telomerase. Embo j. 2006;25(5):1148–59.

36. Oganesian L, Graham ME, Robinson PJ, Bryan TM. Telomerase recognizes G-quadruplex and linear DNA as distinct substrates. Biochemistry. 2007;46(40):11279–90.

37. Lee HT, Bose A, Lee CY, Opresko PL, Myong S. Molecular mechanisms by which oxidative DNA damage promotes telomerase activity. Nucleic Acids Res. 2017;45(20):11752–65.

38. Palacky J, Vorlickova M, Kejnovska I, Mojzes P. Polymorphism of human telomeric quadruplex structure controlled by DNA concentration: a Raman study. Nucleic Acids Res. 2013;41(2):1005–16.

39. Petraccone L, Malafronte A, Amato J, Giancola C. G-quadruplexes from human telomeric DNA: how many conformations in PEG containing solutions? J Phys Chem B. 2012;116(7):2294–305.

40. Long X, Stone MD. Kinetic partitioning modulates human telomere DNA G-quadruplex structural polymorphism. PLoS One. 2013;8(12):e83420.

41. Dai J, Carver M, Punchihewa C, Jones RA, Yang D. Structure of the Hybrid-2 type intramolecular human telomeric G-quadruplex in K+ solution: insights into structure polymorphism of the human telomeric sequence. Nucleic Acids Res. 2007;35(15):4927–40.

42. Peng CG, Damha MJ. G-quadruplex induced stabilization by 2’-deoxy-2’-fluoro-D-arabinonucleic acids (2’F-ANA). Nucleic Acids Res. 2007;35(15):4977–88.

43. Lim KW, Amrane S, Bouaziz S, Xu W, Mu Y, Patel DJ, … Phan AT. Structure of the human telomere in K+ solution: a stable basket-type G-quadruplex with only two G-tetrad layers. Journal of the American Chemical Society. 2009;131(12):4301–9.

44. Abou Assi H, El-Khoury R, Gonzalez C, Damha MJ. 2’-Fluoroarabinonucleic acid modification traps G-quadruplex and i-motif structures in human telomeric DNA. Nucleic Acids Res. 2017;45(20):11535–46.

45. Byrd AK, Raney KD. A parallel quadruplex DNA is bound tightly but unfolded slowly by pif1 helicase. J Biol Chem. 2015;290(10):6482–94.

46. Gavathiotis E, Searle MS. Structure of the parallel-stranded DNA quadruplex d(TTAGGGT)4 containing the human telomeric repeat: evidence for A-tetrad formation from NMR and molecular dynamics simulations. Org Biomol Chem. 2003;1:1650–6.

47. Lingner J, Hughes TR, Shevchenko A, Mann M, Lundblad V, Cech TR. Reverse transcriptase motifs in the catalytic subunit of telomerase. Science. 1997;276(5312):561–7.

48. Harrington L, Zhou W, McPhail T, Oulton R, Yeung DS, Mar V, … Robinson MO. Human telomerase contains evolutionarily conserved catalytic and structural subunits. Genes Dev. 1997;11(23):3109–15.

49. Weinrich SL, Pruzan R, Ma L, Ouellette M, Tesmer VM, Holt SE, … Morin GB. Reconstitution of human telomerase with the template RNA component hTR and the catalytic protein subunit hTRT. Nat Genet. 1997;17(4):498–502.

50. Counter CM, Meyerson M, Eaton EN, Weinberg RA. The catalytic subunit of yeast telomerase. Proc Natl Acad Sci U S A. 1997;94(17):9202–7.

51. Wyatt HD, Lobb DA, Beattie TL. Characterization of physical and functional anchor site interactions in human telomerase. Mol Cell Biol. 2007;27(8):3226–40.

52. Arthanari H, Basu S, Kawano TL, Bolton PH. Fluorescent dyes specific for quadruplex DNA. Nucleic Acids Res. 1998;26(16):3724–8.

53. Nicoludis JM, Barrett SP, Mergny JL, Yatsunyk LA. Interaction of human telomeric DNA with N-methyl mesoporphyrin IX. Nucleic Acids Res. 2012;40(12):5432–47.

54. Samosorn S, Tanwirat B, Muhamad N, Casadei G, Tomkiewicz D, Lewis K, … Bremner JB. Antibacterial activity of berberine-NorA pump inhibitor hybrids with a methylene ether linking group. Bioorg Med Chem. 2009;17(11):3866–72.

55. Samosorn S, inventor13-Arylalkyleneoxyberberine Deriavatives as Anti-Breast Cancer Agents. Thailand 2016.

56. De Cian A, Delemos E, Mergny JL, Teulade-Fichou MP, Monchaud D. Highly efficient G-quadruplex recognition by bisquinolinium compounds. Journal of the American Chemical Society. 2007;129(7):1856–7.

57. Chung WJ, Heddi B, Hamon F, Teulade-Fichou MP, Phan AT. Solution structure of a G-quadruplex bound to the bisquinolinium compound Phen-DC(3). Angew Chem Int Ed Engl. 2014;53(4):999–1002.

58. Ghosh S, Pradhan SK, Kar A, Chowdhury S, Dasgupta D. Molecular basis of recognition of quadruplexes human telomere and c-myc promoter by the putative anticancer agent sanguinarine. Biochim Biophys Acta. 2013;1830(8):4189–201.

59. Gray DM, Wen JD, Gray CW, Repges R, Repges C, Raabe G, Fleischhauer J. Measured and calculated CD spectra of G-quartets stacked with the same or opposite polarities. Chirality. 2008;20(3-4):431–40.

60. Hwang H, Kreig A, Calvert J, Lormand J, Kwon Y, Daley JM, … Myong S. Telomeric overhang length determines structural dynamics and accessibility to telomerase and ALT-associated proteins. Structure. 2014;22(6):842–53.

61. Wang F, Tang ML, Zeng ZX, Wu RY, Xue Y, Hao YH, … Tan Z. Telomere- and telomerase-interacting protein that unfolds telomere G-quadruplex and promotes telomere extension in mammalian cells. Proc Natl Acad Sci U S A. 2012;109(50):20413–8.

62. Paeschke K, Juranek S, Simonsson T, Hempel A, Rhodes D, Lipps HJ. Telomerase recruitment by the telomere end binding protein-beta facilitates G-quadruplex DNA unfolding in ciliates. Nat Struct Mol Biol. 2008;15(6):598–604.

63. Postberg J, Tsytlonok M, Sparvoli D, Rhodes D, Lipps HJ. A telomerase-associated RecQ protein-like helicase resolves telomeric G-quadruplex structures during replication. Gene. 2012;497(2):147–54.

64. Birrento ML, Bryan TM, Samosorn S, Beck JL. ESI-MS Investigation of an Equilibrium between a Bimolecular Quadruplex DNA and a Duplex DNA/RNA Hybrid. J Am Soc Mass Spectrom. 2015;26(7):1165–73.

65. Chen MC, Tippana R, Demeshkina NA, Murat P, Balasubramanian S, Myong S, Ferre-D’Amare AR. Structural basis of G-quadruplex unfolding by the DEAH/RHA helicase DHX36. Nature. 2018;558(7710):465–9.

66. Ray S, Qureshi MH, Malcolm DW, Budhathoki JB, Celik U, Balci H. RPA-mediated unfolding of systematically varying G-quadruplex structures. Biophys J. 2013;104(10):2235–45.

67. Qi X, Xie M, Brown AF, Bley CJ, Podlevsky JD, Chen JJ. RNA/DNA hybrid binding affinity determines telomerase template-translocation efficiency. Embo j. 2012;31(1):150–61.

68. Lacroix L, Seosse A, Mergny JL. Fluorescence-based duplex-quadruplex competition test to screen for telomerase RNA quadruplex ligands. Nucleic Acids Res. 2011;39(4):e21.

69. Nicoludis JM, Miller ST, Jeffrey PD, Barrett SP, Rablen PR, Lawton TJ, Yatsunyk LA. Optimized end-stacking provides specificity of N-methyl mesoporphyrin IX for human telomeric G-quadruplex DNA. Journal of the American Chemical Society. 2012;134(50):20446–56.

70. Gueddouda NM, Mendoza O, Gomez D, Bourdoncle A, Mergny JL. G-quadruplexes unfolding by RHAU helicase. Biochim Biophys Acta Gen Subj. 2017;1861(5 Pt B):1382–8.

71. Schaffitzel C, Berger I, Postberg J, Hanes J, Lipps HJ, Pluckthun A. In vitro generated antibodies specific for telomeric guanine-quadruplex DNA react with Stylonychia lemnae macronuclei. Proc Natl Acad Sci U S A. 2001;98(15):8572–7.

72. Paeschke K, Simonsson T, Postberg J, Rhodes D, Lipps HJ. Telomere end-binding proteins control the formation of G-quadruplex DNA structures in vivo. Nat Struct Mol Biol. 2005;12(10):847–54.

73. Zhang ML, Tong XJ, Fu XH, Zhou BO, Wang J, Liao XH, … Zhou JQ. Yeast telomerase subunit Est1p has guanine quadruplex-promoting activity that is required for telomere elongation. Nat Struct Mol Biol. 2010;17(2):202–9.

74. Murat P, Balasubramanian S. Existence and consequences of G-quadruplex structures in DNA. Curr Opin Genet Dev. 2014;25:22–9.

75. Bao HL, Liu HS, Xu Y. Hybrid-type and two-tetrad antiparallel telomere DNA G-quadruplex structures in living human cells. Nucleic Acids Res. 2019;47(10):4940–7.

76. Liu HY, Zhao Q, Zhang TP, Wu Y, Xiong YX, Wang SK, … Huang ZS. Conformation Selective Antibody Enables Genome Profiling and Leads to Discovery of Parallel G-Quadruplex in Human Telomeres. Cell chemical biology. 2016;23(10):1261–70.

77. Sen D, Gilbert W. Formation of parallel four-stranded complexes by guanine-rich motifs in DNA and its implications for meiosis. Nature. 1988;334(6180):364–6.

78. Tomlinson CG, Sasaki N, Jurczyluk J, Bryan TM, Cohen SB. Quantitative assays for measuring human telomerase activity and DNA binding properties. Methods. 2017;114:85–95.

79. Cohen SB, Reddel RR. A sensitive direct human telomerase activity assay. Nat Methods. 2008;5(4):355–60.

