## Supplementary table and figures for "A mechanism for the extension and unfolding of parallel telomeric G-quadruplexes by human telomerase at single-molecule resolution"

**Supplementary Table 1:** Oligonucleotides used in this study

| **Name** | **Sequence (5' to 3')** | **Extinction coefficient (L*M$^{-1}$cm$^{-1}$)** |
|---|---|---|
| 22G0 | AGGGTTAGGGTTAGGGTTAGGG | 228,500 |
| 22G3 | A(fG)GGTTA(fG)(fG)GTTA(fG)(fG)GTTA(fG)GG | 228,500 |
| 22G0+tail | TGGCGACGGCAGCGAGGCTAGGGTTAGGGTTAGGGTTAGGG | 407,300 |
| 22G3+tail | TGGCGACGGCAGCGAGGCTA(fG)GGTTA(fG)(fG)GTTA(fG)(fG)G T$^{AlexaFluor555}$TA(fG)GG | 407,300 |
| 647-Strand2 | GCCTCGCT$^{AlexaFluor647}$GCCGTCGCCA-Spacer18-Biotin | 192,600 |
| 7GGT | TTAGGGT | 69,800 |
| Bio-sp-7GGT | Biotin-Spacer18-TTAGGGT | 69,800 |
| 647-7GGT | Spacer18-TT$^{AlexaFluor647}$AGGGT | 69,800 |
| 555-7GGT | Spacer18-TT$^{AlexaFluor555}$AGGGT | 69,800 |
| Bio-L-18GGG | Biotin-CTAGACCTGTCATCATTAGGGTTAGGGTTAGGG | 325,000 |
| RNA.10C | CUAACCCUAA | 98,200 |
| DNA.10C | CTAACCCTAA | 96,600 |

(fG) = 2'F-araG

### Supplementary Figure 1

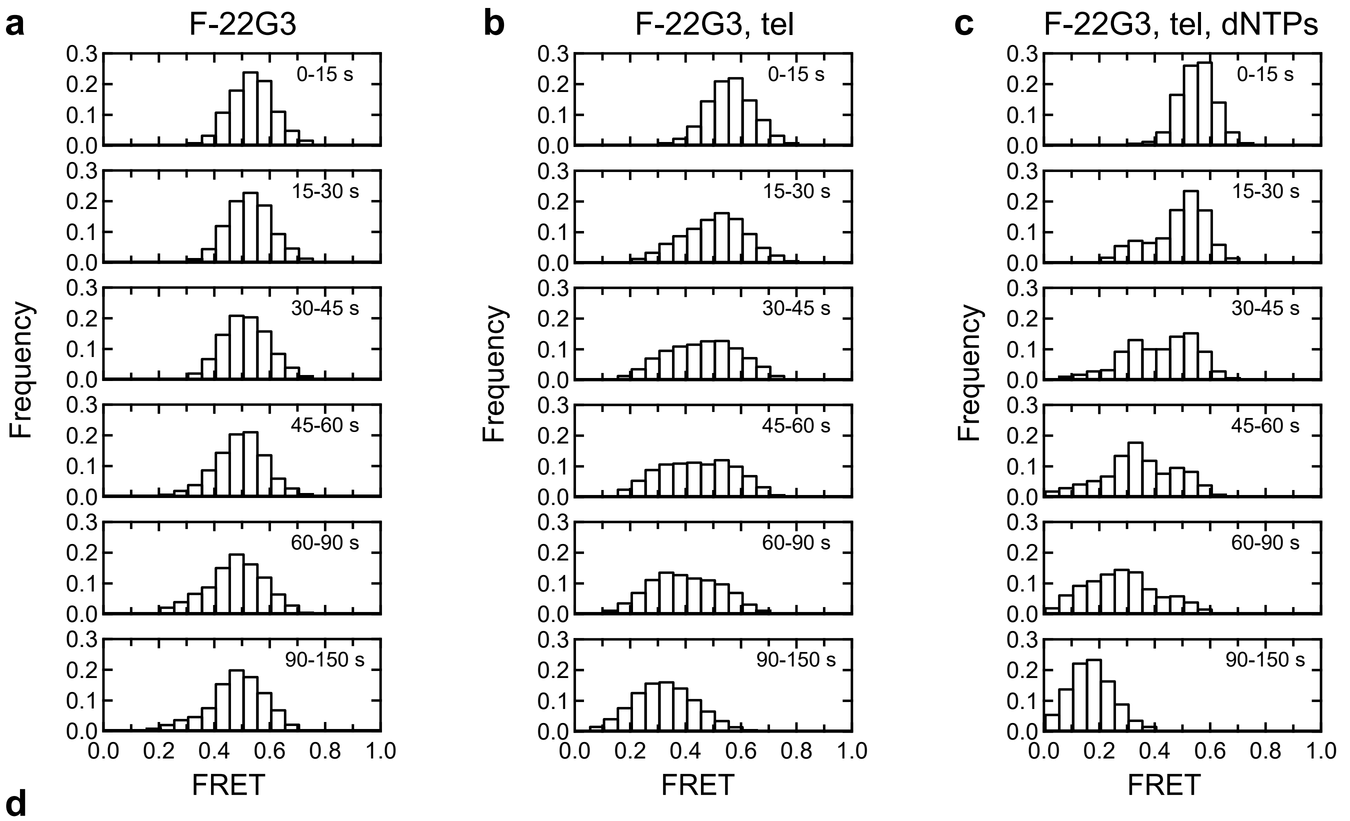


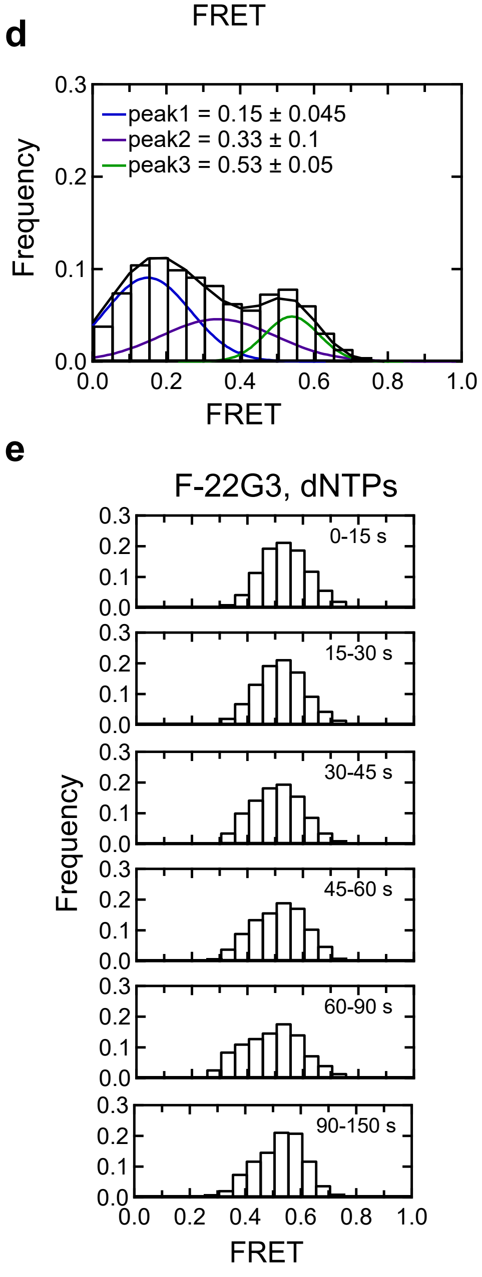


**Supplementary Figure 1 (related to Figure 1): Telomerase binds, extends and unfolds a parallel intramolecular G-quadruplex in the presence of dNTPs.**

**(a),(b),(c),(e)** Histograms showing changes in FRET values of a population of F-22G3 molecules under different experimental conditions over 150 s, binned into 15 - 60 s time intervals.

**(d)** Histogram showing three different FRET populations in the presence of telomerase and dNTPs; a multiple Gaussian fit was applied to obtain the mean FRET peaks of these populations, with errors representing the curve fitting error.

n = 99 **(a)**, 125 **(b)**, 90 **(c)**,**(d)**, and 44 **(e)** molecules, collected in 2 - 4 independent experiments.

### Supplementary Figure 2

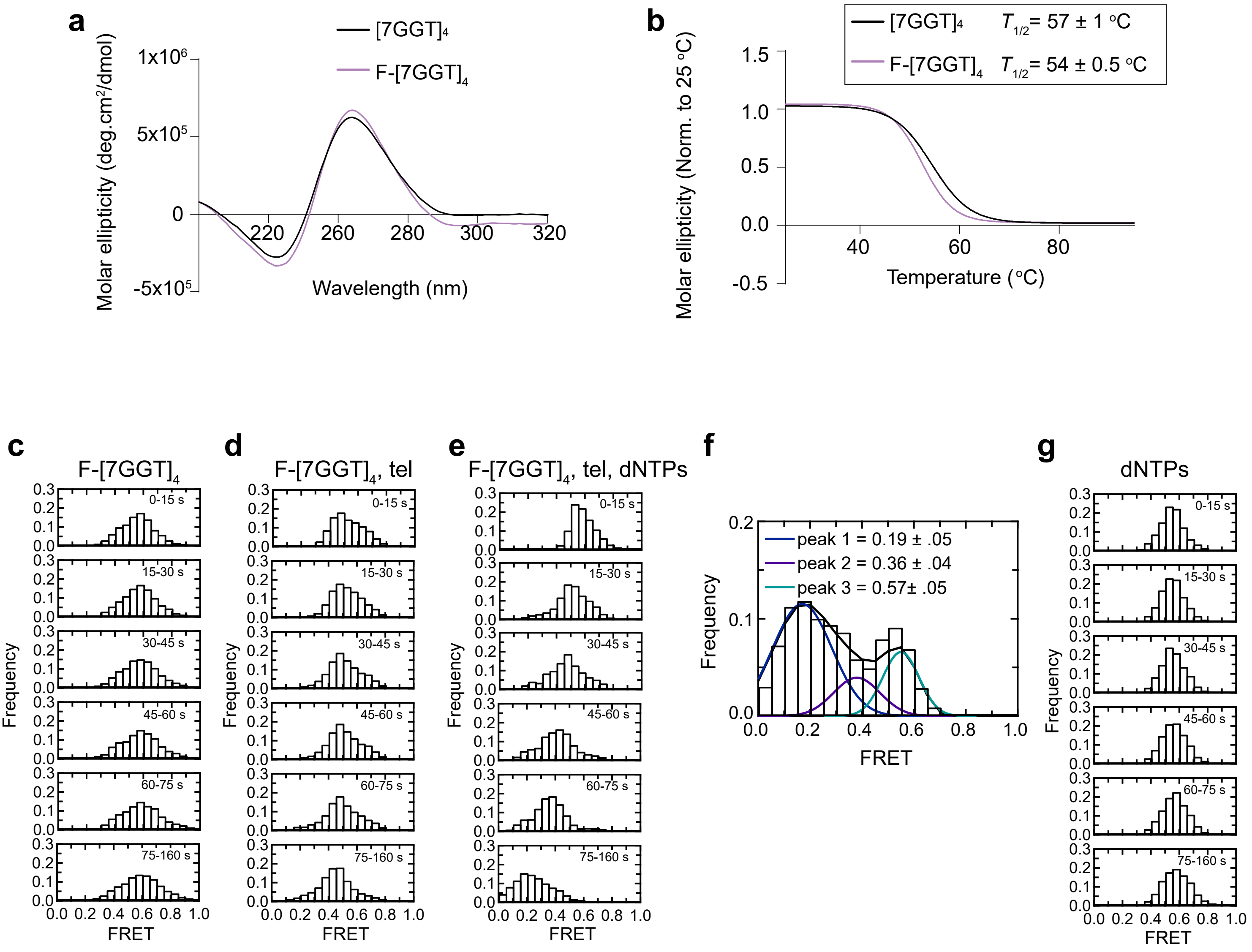


**Supplementary Figure 2 (related to Figure 2): Telomerase binds, extends and unfolds a stable parallel tetrameric G-quadruplex.** (**a)** Circular dichroism (CD) spectra of 10 μM fluorophore-modified F-$[7GGT]_4$ and unmodified $[7GGT]_4$. (**b)** Thermal stability of modified and unmodified $[7GGT]_4$, measured using CD at 260 nm. Since intermolecular G-quadruplexes reform very slowly and are therefore not at equilibrium during the measurement, we refer to this value as $T_{½}$ rather than $T_m$, which is the true thermodynamic parameter at equilibrium. Mean ± SD of 3 independent experiments. **(c),(d),(e),(g)** Histograms showing changes in FRET values of a population of molecules under different experimental conditions over 160 s, binned into 15 s time intervals. **(f)** Histogram showing three different FRET populations in the presence of telomerase and dNTPs; a multiple Gaussian fit was applied to obtain the mean FRET peaks of these populations, with errors representing the curve fitting error. n = 105 **(c)**, 87 **(d)**, 81 **(e)**,**(f)**, and 110 **(g)** molecules, collected in 4 – 6 independent experiments.

#### Supplementary Figure 3

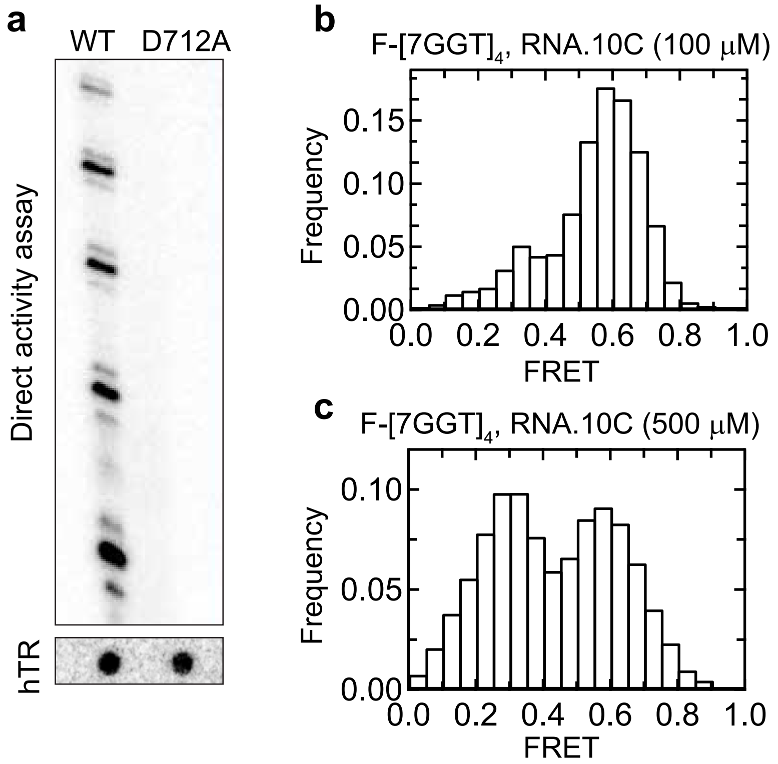


**Supplementary Figure 3 (related to Figure 3): (a)** Telomerase activity assay using purified catalytically inactive human telomerase (D712A) compared with wild type (WT) telomerase. Upper gel shows products of telomerase extension of oligonucleotide Bio-L-18GGG (1 μM; Supplementary Table 1); lower panel shows a northern blot of the amount of hTR in each telomerase preparation, to control for equal amounts of enzyme. **(b), (c)** Histograms of the FRET values of a population of F-$[7GGT]_4$ molecules, in the presence of the indicated concentrations of oligonucleotide RNA.10C; n = 84 (**b**) and 129 (**c**) molecules, collected in 4 – 6 independent experiments.

#### Supplementary Figure 4

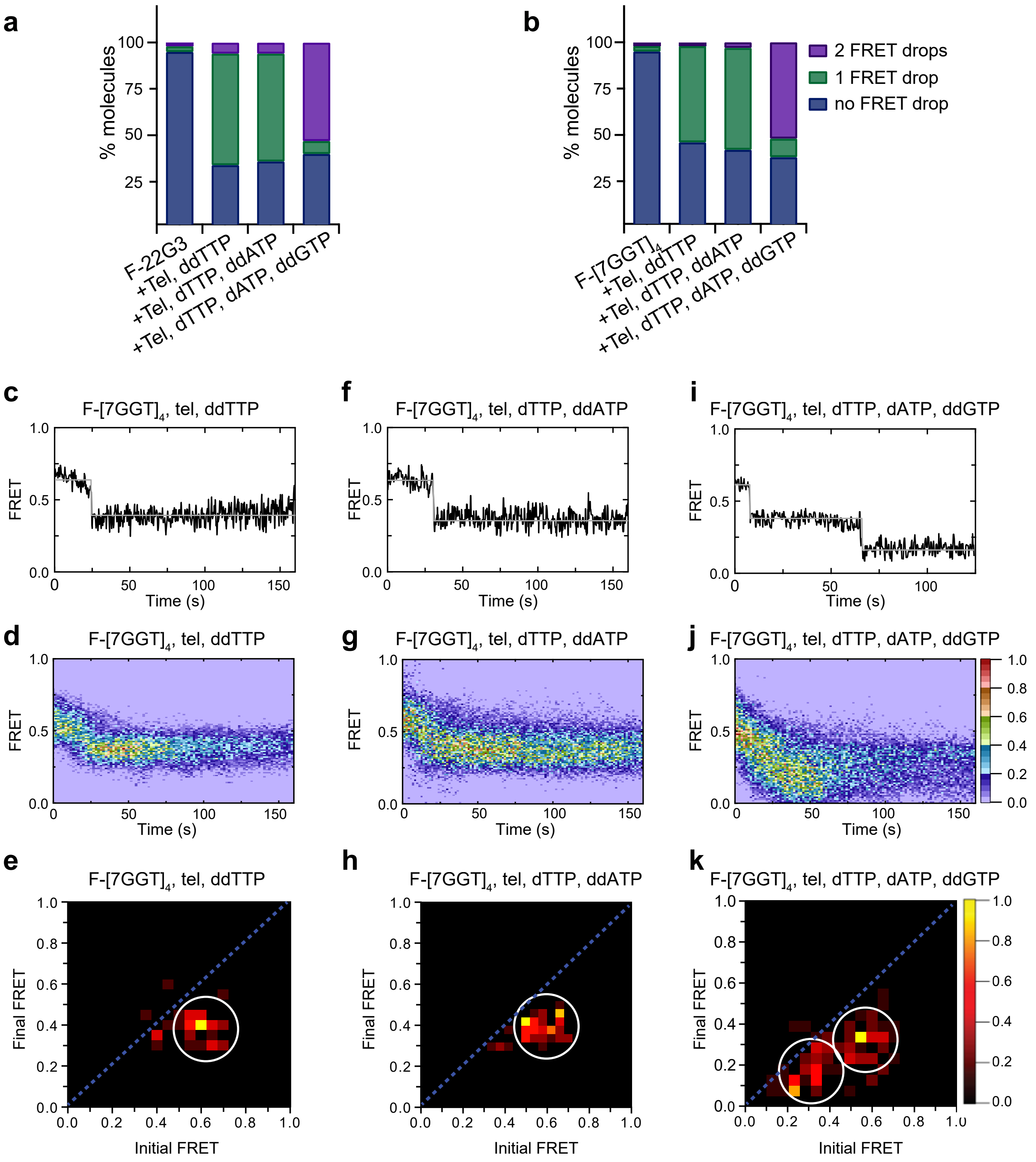


**Supplementary Figure 4 (related to Figure 4): Telomerase unfolds intra- and intermolecular G4 using same mechanism. (a - b)** Plot of the percentage of molecules showing no change in FRET, a single FRET drop, or a two-step FRET drop, during telomerase extension of F-22G3 **(a)** or F-$[7GGT]_4$ **(b)**. **(c),(f),(i)** Examples of individual FRET trajectories in the presence of telomerase and the indicated combinations of nucleotides. **(d),(g),(j)** Heat maps of the distribution of FRET intensities over 0 - 150 s in 69, 73 and 80 molecules, respectively, in the presence of telomerase and the indicated combinations of nucleotides. All plots include molecules collected in 4 – 6 independent experiments. For color key, see panel **(j)**. **(e),(h),(k)** TDPs showing the changes between the initial and final FRET values of F-$[7GGT]_4$ in the presence of telomerase and the indicated combinations of nucleotides (n = 69, 73 and 80 molecules, respectively). For color key, see panel **(k)**.

#### Supplementary Figure 5

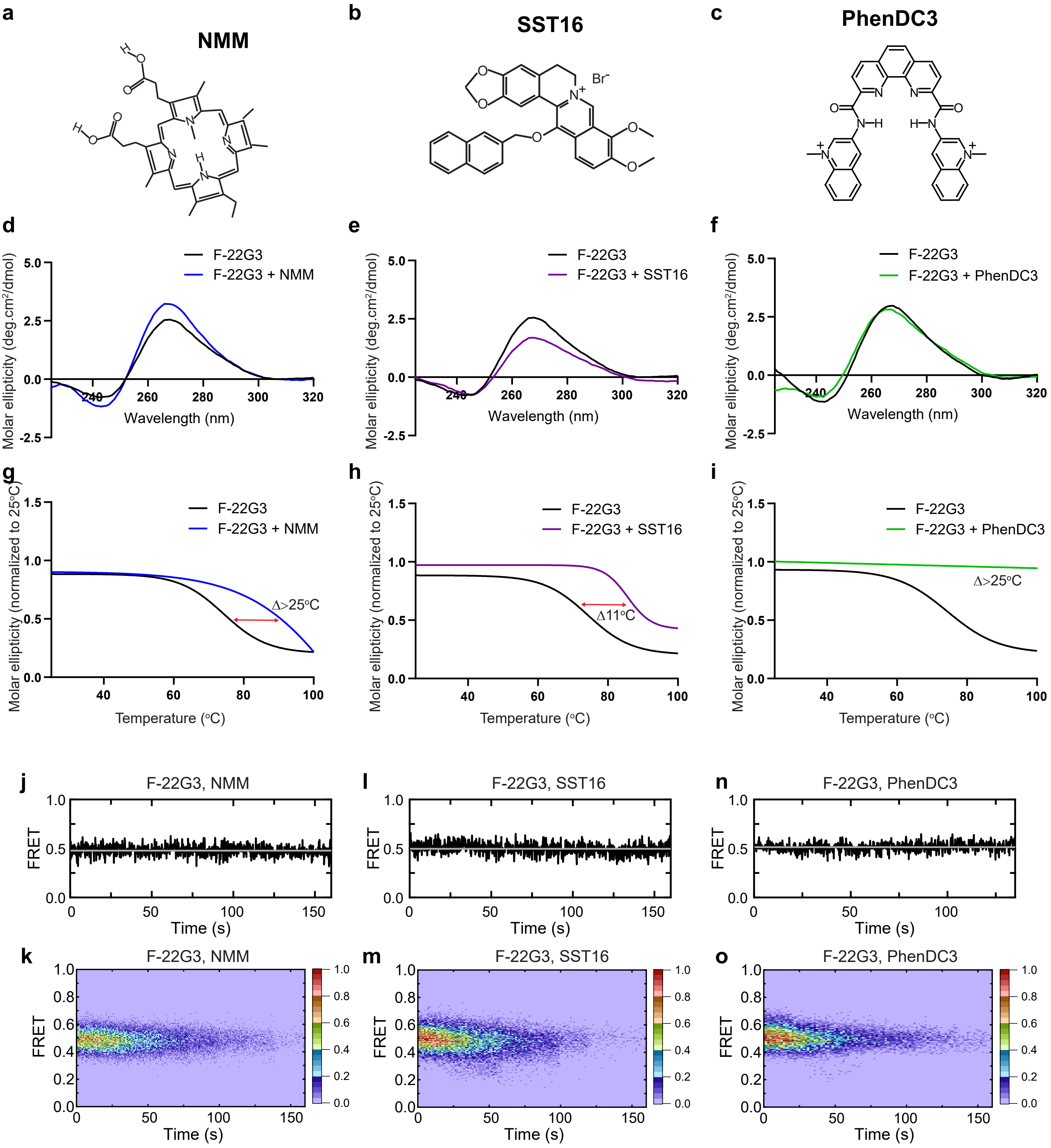


**Supplementary Figure 5 (related to Figure 5): Stabilization of parallel intramolecular G4 F-22G3 by ligands NMM, SST16 and PhenDC3. (a - c)** Structures of NMM, SST16 and PhenDC3. **(d - i)** CD spectra and melting curves measured by CD at 260 nm of 250 nM F-22G3 in the presence or absence of 40 µM NMM, 5 µM SST16 or 1 µM PhenDC3. NMM was incubated with F-22G3 during G4 folding at 20-fold higher concentrations, and the G4-ligand complex diluted for CD measurement. **(j - o)** Representative FRET trajectories and heat maps of F-22G3 in the presence of NMM (800 µM in folding reaction), SST16 (5 µM) or PhenDC3 (1 µM); n = 110 (**k**), n = 96 (**m**), n = 103 (**o**), collected in 2-3 independent experiments.

#### Supplementary Figure 6

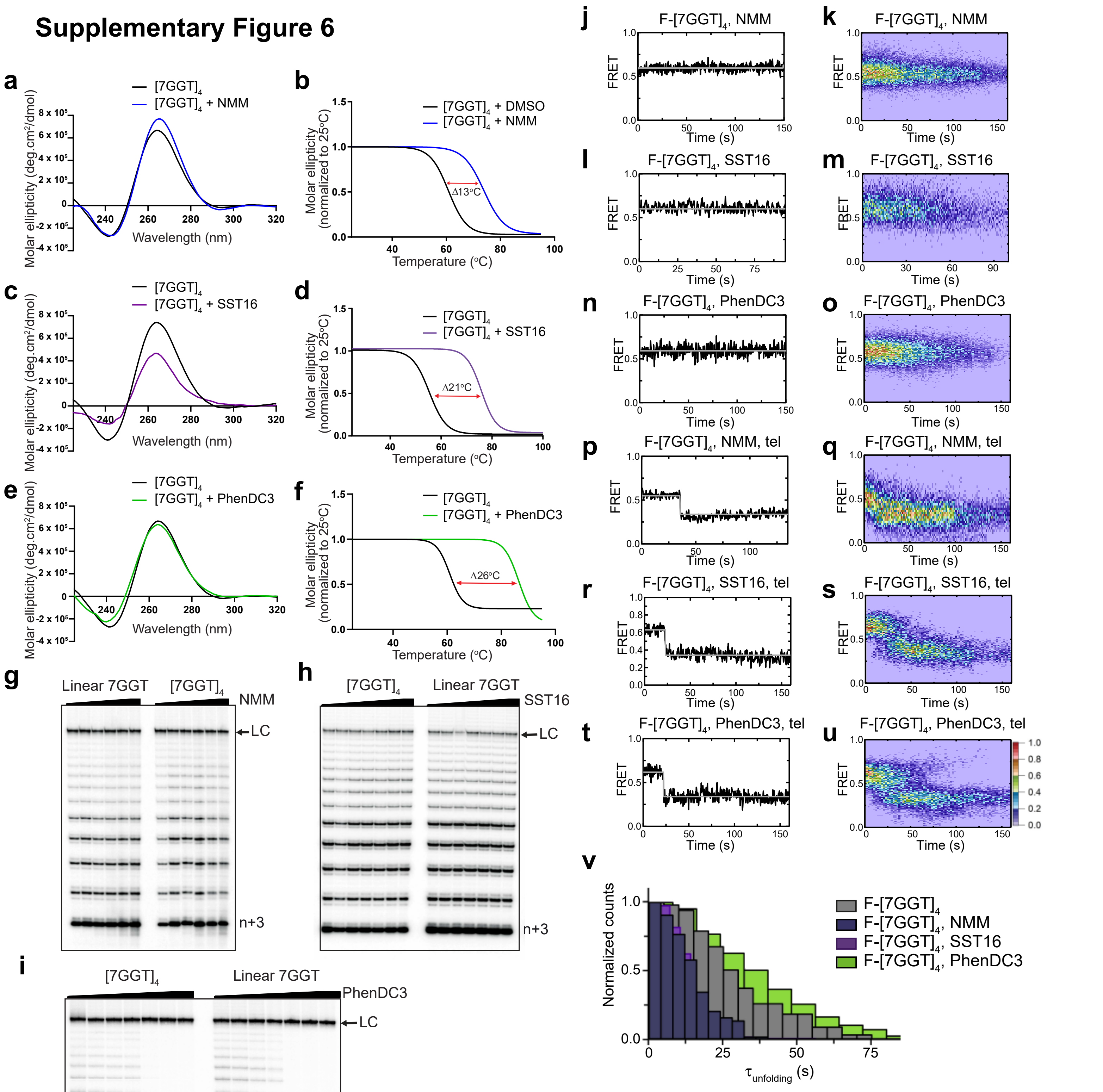


**Supplementary Figure 6 (related to Figure 6): Telomerase extension of $[7GGT]_4$ is not inhibited by the binding of small-molecule ligands NMM, SST16 and PhenDC3.**

**(a - f)** CD spectra and melting curves measured by CD at 260 nm of 1 μM $[7GGT]_4$ in the presence or absence of 40 μM NMM, 100 μM SST16 or 1 μM PhenDC3. **(g - i)** Telomerase extension assays in the presence of either NMM (0, 4 μM, 20 μM, 40 μM, 80 μM, 160 μM), SST16 (0, 10 μM, 25 μM, 50 μM, 75 μM, 100 μM) or PhenDC3 (0, 0, 0.5 μM, 1 μM, 2.5 μM, 5 μM, 10 μM). DNA concentrations were 1 μM (**g**, **h**) or 250 nM (**i**). NMM was incubated with 7GGT during G4 folding at 250-fold higher concentrations, and the G4-ligand complex diluted for the assay. LC is a 100 nt loading control and n+3 indicates the product with the first three nucleotides incorporated. **(j - u)** Representative FRET trajectories and heat maps of F-$[7GGT]_4$ in the presence of NMM (10 mM in folding reaction), SST16 (100 μM) or PhenDC3 (1 μM), with (**p** - **u**) or without (**j** - **o**) telomerase; n = 120 (**k**), 73 (**m**), 111 (**o**), 80 (**q**), 60 (**s**), 74 (**u**) molecules, collected in 4 – 6 independent experiments. For color key, see panel (**u**). **(v)** The unfolding rate of F-$[7GGT]_4$ in the presence of NMM, SST16, PhenDC3 or no ligand, calculated by fitting the dwell time distributions of the intermediate FRET state to a single exponential equation. n = 55 (no ligand), 44 (NMM), 60 (SST16) and 71 (PhenDC3).
